# Formation of uniform reaction volumes using concentric amphiphilic microparticles

**DOI:** 10.1101/2020.03.15.992321

**Authors:** Ghulam Destgeer, Mengxing Ouyang, Chueh-Yu Wu, Dino Di Carlo

## Abstract

Reactions performed in uniform microscale volumes have enabled numerous applications in the analysis of rare entities (e.g. cells and molecules), however, sophisticated instruments are usually required to form large numbers of uniform compartments. Here, uniform aqueous droplets are formed by simply mixing microscale multi-material particles, consisting of concentric hydrophobic outer and hydrophilic inner layers, with oil and water. The particles are manufactured in batch using a 3D printed device to co-flow four concentric streams of polymer precursors which are polymerized with UV light. The size of the particles is readily controlled by adjusting the fluid flow rate ratios and mask design; whereas the cross-sectional shapes are altered by microfluidic nozzle design in the 3D printed device. Once a particle encapsulates an aqueous volume, each “dropicle” provides uniform compartmentalization and customizable shape-coding for each sample volume to enable multiplexing of uniform reactions in a scalable manner. We implement an enzymatically-amplified affinity assay using the dropicle system, yielding a detection limit of <1 pM with a dynamic range of at least 3 orders of magnitude. Moreover, multiplexing using two types of shape-coded particles was demonstrated without cross talk, laying a foundation for democratized single-entity assays.

## 1. Introduction

Breaking a sample volume into numerous small compartments enables the accumulation of signal from a small number of molecules or cells to detectable levels in a reasonable time period. Uniformity in the compartment volumes ensures the reaction conditions are relatively similar and reactions across volumes can be compared. Microfluidic wells^[1–3]^ or droplet generators^[4–6]^ enable uniform compartmentalization of a sample fluid volume into many smaller reactions, however skilled users or specialized and costly commercial instruments have been required for reproducible implementation. In addition, for many affinity assays, a solid phase, such as microbeads, is desired to bind a target and allow washing to remove excess reagents or minimize non-specific binding. A solid phase also allows barcoding for multiplex detection, either by the shape^[7–9]^ and/or color^[10–12]^ and binding and growth of adherent cells. However, introducing a microbead into a well or droplet may introduce other challenges associated with uniform loading of compartments.

Droplet microfluidics is a commonly used compartmentalization technique to disperse aqueous assay reagents with cells or molecules into small segmented volumes inside a continuous oil phase for subsequent signal amplification and detection without cross talk.^[5]^ However, encapsulating a single microbead as a solid phase inside each droplet is limited by Poisson loading statistics.^[13]^ Therefore, the success rate for encapsulating a combination of exactly two distinct components i.e. a microbead and a target cell or molecule inside a droplet approaches ~1% of the entire population of droplets, whereas the remaining droplets would have undesired combinations of the components.^[4,14,15]^ Moreover, for multiplexing, multiple types of microbeads with distinct barcoding signatures should be encapsulated in separate droplets with the targets of interest, which is further limited by multiplicative probabilities to triple or larger Poisson distributions for duplex or greater multiplexing.

Therefore, an instrument-free compartmentalization system that forms uniform droplets embedded with a single solid phase per droplet can address many of the challenges with current approaches.

An attractive alternative technique to create homogeneous compartments uses uniform engineered microparticles to form aqueous volumes by simple exchange of fluids. However, monolithic creation of particles that have structures and material chemistries tuned to spontaneously collect defined uniform aqueous volumes in an oil continuous phase is challenging, and has not yet been reported to our knowledge. Spherical gel beads have been vigorously mixed with aqueous solutions and chemical surfactants to create aqueous volumes occupying a thin shell around the bead, however uniform drops are not at an interfacial energy minimum of the system, while the absence of a cavity has precluded use with mammalian cells and may inhibit some reactions.^[16,17]^ In addition, spherical particles have more limited barcoding options, and it has been reported that the presence of surfactants leads to higher rates of transport of products of reactions through the continuous oil phase, reducing sensitivity of enzymatic assays.^[18,19]^

A range of fabrication methodologies have been explored over the past decade to create particles with different shapes and functionalities using continuous^[20–24]^ or stop flow lithography techniques^[25–28]^ combined with hydrodynamic focusing,^[29–32]^ magnetically tunable color printing,^[12,33]^ vertical flows,^[34]^ structured hollow fibers^[35]^ or inertial forces.^[36–39]^ Particles comprised of layers of hydrophobic and hydrophilic materials were shown to selectively interact and assemble around aqueous drops.^[22,40]^ However, these approaches either do not hold a uniform volume of a compartmentalized aqueous phase^[22]^ or suffer from a low throughput and complicated fabrication workflow (Table S1).^[40]^ In addition, most of these techniques depend on a thin polymerization-inhibition layer close to the walls of oxygen-permeable polydimethylsiloxane (PDMS) microfluidic channels that require cleanroom fabrication facilities. Alternative techniques to shape precursor flows using inertial flow sculpting^[38]^ removes limitations on an oxygen-permeable PDMS layer but consumes more reagents per particle fabricated and requires high-pressure flow, whereas vertical flow lithography^[34]^ and maskless lithography^[40,41]^ that have the capability to create multi-layer 3D particles, suffer from a limited throughput.

Here, we use a 3D printed microfluidic channel network to create particles with an amphiphilic chemistry and a concentric ring-shaped geometry we hypothesized would encompass a uniform volume of aqueous phase inside a hydrophilic cavity of the particle. By using 3D printed concentrically stacked channels, manufactured without cleanroom facilities, we achieve a hydrodynamically focused co-axial flow of hydrophobic poly(propylene glycol) diacrylate (PPGDA) and hydrophilic poly(ethylene glycol) diacrylate (PEGDA) polymers. Co-flowing polymer precursor streams with photo-initiator (PI) are exposed to UV light through a photomask to create concentric particles, whereas an inert outer sheath flow prevents the particles from sticking to the glass capillary walls and an inert inner sheath flow defines the open cavity of the particle. The size of the particle (i.e., 340-400 μm) and the cavity (i.e., 100-200 μm) are readily controlled by adjusting the PPGDA to PEGDA flow rate ratio (i.e., 4:1), whereas the shapes of the particles are modulated based on the 3D printed channel designs.

These multi-material concentric amphiphilic particles in which we further functionalize the inner layer to capture target molecules are shown to spontaneously form uniform aqueous drops with assay reagents upon solution exchange. The inner hydrophilic surface of the particle associates with the aqueous phase while the outer hydrophobic surface prefers the oil phase upon exchange to an oil continuous phase. Given the energetic stability of this configuration, simple transfer steps of aqueous reagents with the amphiphilic particles yields uniform aqueous volumes, without the need for control of drop breakup mechanisms that employ precise control of flow rates or pressures. The aqueous drops contain assay reagents surrounded by a continuous oil phase that prevents cross talk during long-duration reactions, while the solid substrate of the templating particle provides an anchor for capturing target molecules. The need for encapsulating an additional particle inside the droplet is also eliminated, thus reducing the effect of Poisson statistics on encapsulation performance.

We conduct an amplified assay using standard reagents for enzyme linked immunosorbent assays (ELISA) within droplets formed by these amphiphilic particles and achieve a sub-pM detection limit with wide dynamic range by accumulating results from hundreds of parallel reactions. A collection of dropicles, each acting as part of a “swarm” of individual sensors,^[42]^ improves the statistical accuracy in quantitative prediction of concentration by averaging out small differences in reactions across the dropicles. Reactions proceeding simultaneously in two different types of shape-coded particles, separately functionalized, yield minimal cross talk. Tunable assay performance (i.e. detection limit and dynamic range) is also achieved by adjusting the particle dimensions and materials, which is desirable for multiplex detection of biomarkers spanning a large range of clinically relevant concentrations.

## 2. Results

### 2.1. Amphiphilic, Shape-Coded, and Size-Tunable Particle Fabrication

We develop a new approach, leveraging 3D printing, in order to fabricate uniform particles comprising concentric materials of different hydrophobicity. By using a 3D printed microfluidic network of channels, we are able to route four density matched precursor fluids with different chemistries, an inert outer sheath (PPGDA only, or PPG and PI), PPGDA and PI, PEGDA and PI, and an inert inner sheath (PEGDA only, or PEG and PI), through four concentric channels to obtain a co-axial flow structure (**Figure 1A**). The internal structure of the microfluidic channels with a tapered geometry at the exit ensures that the flow stream from channel 4 first co-flows with the flow stream from channel 3, which is subsequently combined with the flow streams from channel 2 and 1. To reduce the effect of diffusion of species between streams, the diameter of the device is gradually reduced to increase the flow velocity as the flow streams are merged together in a sequential manner, resulting in a final Peclet number (*Pe*) of ~5 × 10^5^. The fully developed co-axial flow is briefly stopped to expose the polymer precursors to UV light through a patterned array of windows in a photomask to cure multi-material concentric particles (Figure 1C). The cured particles with a hydrophobic outer PPG layer and hydrophilic inner PEG layer are washed downstream to a collection tube as the flow inside the channels is restarted (Figure 1D). This cycle is repeated automatically to continuously fabricate and collect particles for a desired number of cycles (Figure 1B). The partitioning of fluorescent resorufin into the PEG layer confirms the multi-material composition of the particles (Figure 1E). The reproducible structured co-flow along with automated processing and exposure ensures that each fabrication batch of the particles has a high uniformity in their shape, size and material composition (Figure 1F-G).

**Figure 1.**
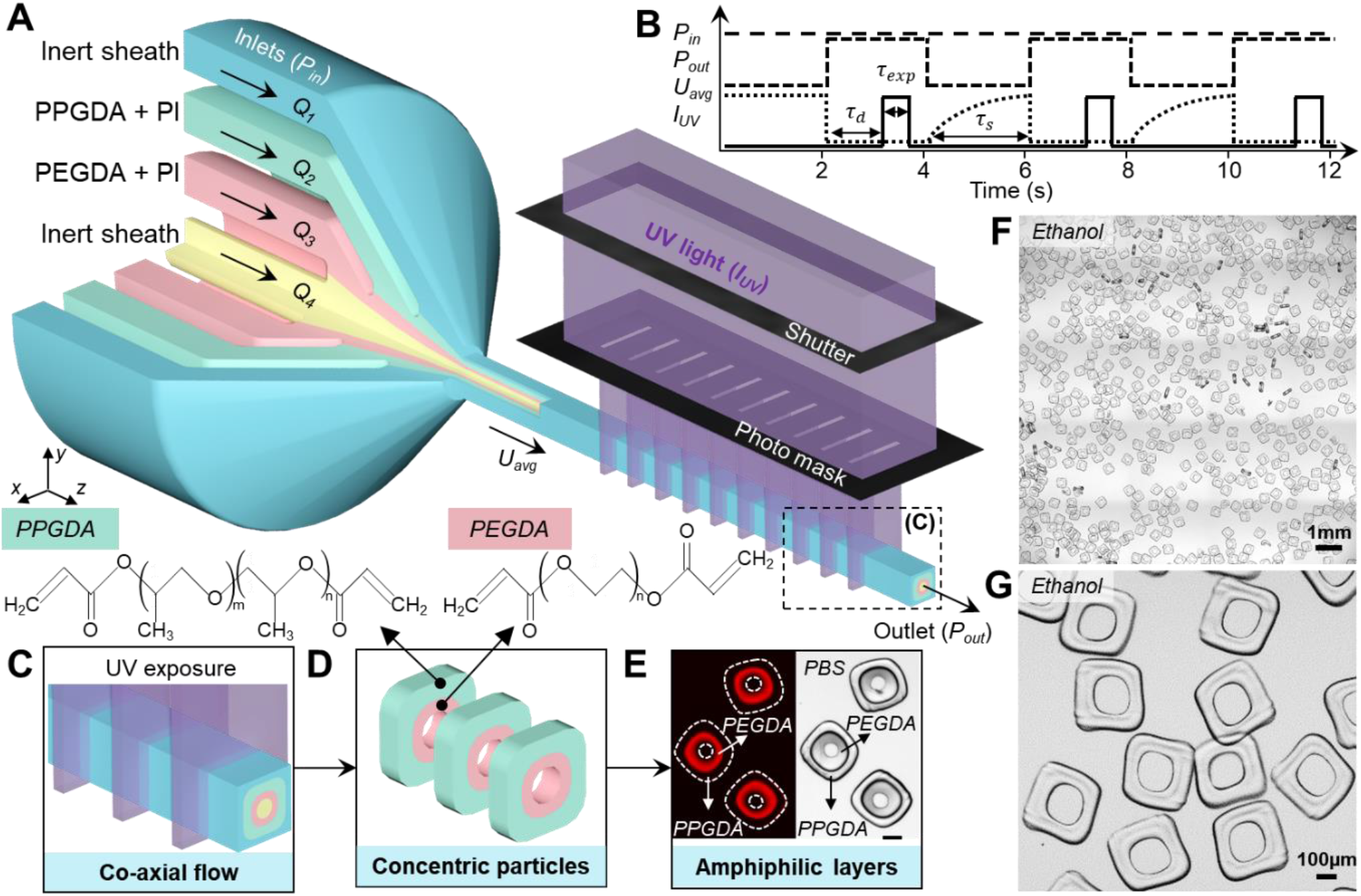
Fabrication of concentric amphiphilic microparticles. **(A)** Four different streams of fluids are pumped through inlets 1-4 at flow rates *Q*_*1*_ to *Q*_*4*_, respectively, resulting in an inlet pressure, *P*_*in*_. Flows from inlets 2 and 3 contain photoinitiator (PI) and polymer pre-cursors. The flow reaches an average velocity of *U*_*avg*_ within the square section of the device. The flow is stopped (*U*_*avg*_ = 0) with a pinch valve resulting in an outlet pressure of *P*_*out*_ (= *P*_*in*_). A shutter is opened after a short delay time (*τ*_*d*_) to expose the flow streams to UV light with intensity *I*_*UV*_ through a photomask for an even smaller exposure time (*τ*_*exp*_). At this point, the pinch valve is opened (*P*_*out*_ = 0) and the syringe pumps are re-started to reach an average flow velocity of *U*_*avg*_ within a flow stabilization time (*τ*_*s*_). **(B)** The plot summarizes the whole process of flow stoppage, UV exposure and flow stabilization in a cyclic manner. **(C)** A zoomed-in view of the exposure region shows that the co-axial streams are exposed to rectangular-shaped UV beams. **(D)** The flow streams containing PI originating from channels 2 and 3 are polymerized in the form of a ring-shaped amphiphilic particle with hydrophobic outer layer made of PPGDA and hydrophilic inner layer made of PEGDA. **(E)** TRITC (left) and bright-field (right) images of representative fabricated particles mixed with resorufin that partitions into the PEG layer are shown. The dashed lines mark the outer boundaries of the particle in the fluorescent image. Scale bar is 100 μm. **(F)** A batch of fabricated O-shaped particles suspended in ethanol shows a high uniformity in the particle size as well as the inner cavity. Scale bar is 1 mm. **(G)** A zoomed image of the particles suspended in ethanol shows a circular cavity within an outer square-shaped boundary of the particle.

Instead of relying on changes to the masked light intersecting the polymer precursor stream we engineer the 3D printed channel structures to tune the cross-sectional flow shapes and produce a variety of shape-coded particles (**Figure 2**). Microfluidic devices that have nozzles designed with different cross-sections (*a-a’*), as indicated in Figure S1A, yield particles with engineered shapes. Particles with shape codes defined in the outer, inner, or both materials are fabricated to demonstrate the capabilities of the 3D printed channels (Figure 2, Figure S1B, Figure S2). For outer shape codes, the four corners of square particles are systematically removed to obtain six different shapes while the inner cavity shape is kept the same (Figure 2C). For inner shape codes, the outer square boundary of particles is maintained, while different distributions of the PEG layer are shaped to form unique internal features (Figure 2D). The outer and inner boundaries of particles are also modified in combination to obtain complex internal and external particle features (Figure 2F). The cross-sectional geometries of shaped particles possess high uniformity with a CV of less than 4% in outer diameter (Figure S1C). The flow rates for the polymer precursors are also readily adjusted to tune the size of the particles (Figure S1D, Figure S3). For O-shaped particles, increasing the flow rate ratio (*Q*_*1,2*_:*Q*_*3,4*_) from 1 to 4 gradually reduced the size of the particle cavities from ~185 μm to ~100 μm (Figure S1D).

**Figure 2.**
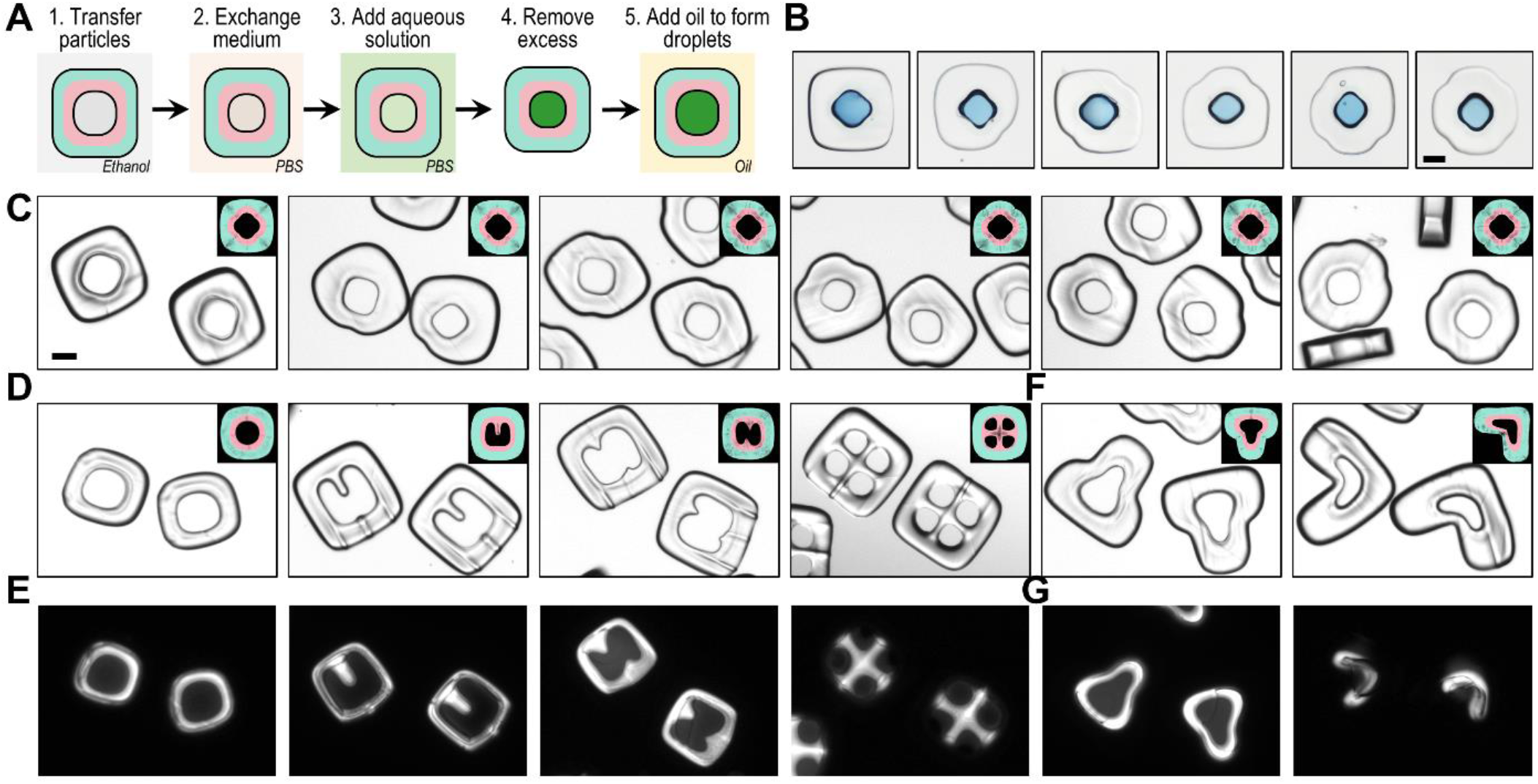
Shape-coded particles and aqueous droplet formation inside the cavities. **(A)** Workflow for dropicle formation as amphiphilic particles are transferred from ethanol to PBS before exchanging with an oil continuous phase. **(B)** Droplet formation using six different outer shape coded particles. **(C)** Outer shape code demonstrated by systematically removing combinations of corners from an outer square shaped particle. **(D)** Inner shape coded particles with an outer square shaped boundary. **(E)** Droplet formation inside inner shape coded particles as observed with fluorescence imaging of encapsulated resorufin dye. Partitioning into the inner PEG layer is observed **(F)** Bilayer-shape coded particles with changes reflected in the inner and outer boundaries of the particles. **(G)** Droplet formation inside bilayer-shape coded particles as observed with fluorescence imaging of encapsulated resorufin dye. Scale bars are 100 μm. Insets for (C, D, F) show fluid dynamic simulation results of the cross-sectional shape of the outer PPG layer (cyan) and inner PEG layer (magenta).

### 2.2. Dropicle Formation

Dropicles, uniformly-sized droplets supported by particles, are formed by simple pipetting for fluid phase exchange (Figure 2A). The surrounding fluid phase for the particles is exchanged first from ethanol to phosphate buffered saline (PBS), and then from PBS to a mixture of PBS with aqueous solution (e.g., a fluorophore or color dye solution) for subsequent intensity measurements or droplet visualization. Adding a final oil phase with low interfacial tension with the outer PPG layer creates hundreds of isolated compartments, or dropicles, immediately. As the excess fluid is removed at each step, nanoliter-scale volumes of aqueous solution remain in an energetically favorable configuration associated with the hydrophilic core of the amphiphilic particles. We would like to point out that the leakage of constituents out of droplets has been a general issue for conventional droplet systems, as these systems require the use of surfactant in the oil phase for droplet formation and stabilization.^[18,19]^ The surfactants can enhance transport between droplets through micellar transport of hydrophilic compounds which may have low partition coefficients into the oil phase. However, it is worth noting that our system does not need surfactant to form droplets. Instead, the droplet formation is due to the hydrophobicity difference within different layers of the multi-material particles. Droplets formed within the various shape coded particles span the entire inner hydrophilic layer and reflect the shape of the cavities (Figure 2B, Figure S1E). Besides manipulating the outer hydrophobic layer of the amphiphilic particles (Figure 2C), variations in the hydrophilic inner cavity are also engineered (Figure 2D, F), however, these inner and outer shape changes do not appear to affect the ability to hold an aqueous droplet (Figure 2E, G). Particles with different designs can hold 2-6 nL droplets depending on the shape and size of the cavities, whereas the variation in volume is less than 10% on average and variation in diameter of an equivalent volume spherical droplet is ~3% (Supporting Information).

### 2.3. Amplified Affinity Assay in Dropicles

The materials and emulsification process used to form dropicles is compatible with affinity assays using enzymatic amplification of signal. We demonstrate a QuantaRed assay within dropicles, in which a fluorogenic precursor (10-Acetyl-3,7-dihydroxyphenoxazine, ADHP) is converted into fluorescent resorufin due to the activity of horse radish peroxidase (HRP) that bound to the surface of biotinylated particles (**Figure 3A**). The assay generally follows standard steps required for conducting ELISAs. Biotinylated particles suspended in ethanol are first added to a well plate with a hydrophobic surface, where they quickly settle on the bottom of the well with the majority facing upward, due to the density difference and aspect ratio of the particles. Next, the ethanol solution is exchanged with a PBS buffer with an average particle retention rate of 98% through solution exchange and subsequent incubation steps (see Supporting Information for details). Then, streptavidin-HRP solution was added and incubated for 30 min, leading to HRP binding to particles. Following HRP binding and washing, QuantaRed solution is loaded into the well with excess removed immediately from the corner of the well. Lastly, oil is added to form and seal the droplets within seconds (see Experimental Section for details). Once in dropicles, ADHP in the QuantaRed solution is catalytically converted by HRP into resorufin which accumulates in the aqueous phase and also partitions to some extent into the encapsulating PEG layer,^[43,44]^ but is not observed to transfer into the oil phase. Fluorescent and bright field images of the dropicles while in oil are obtained for a few hundreds of compartmentalized reactions (Figure 3B). Typically, the fluorescent signal from the droplets increases over time (Figure S4) and as expected increases at a higher rate for higher concentrations of streptavidin-HRP (e.g., 10 pM) while remaining constant for a negative control group of particles that are incubated with PBS only. In further validating this assay, we picked specific time points to image and quantify dropicle fluorescence after which sufficient fluorescent signal is developed (typically > 15 min).

**Figure 3.**
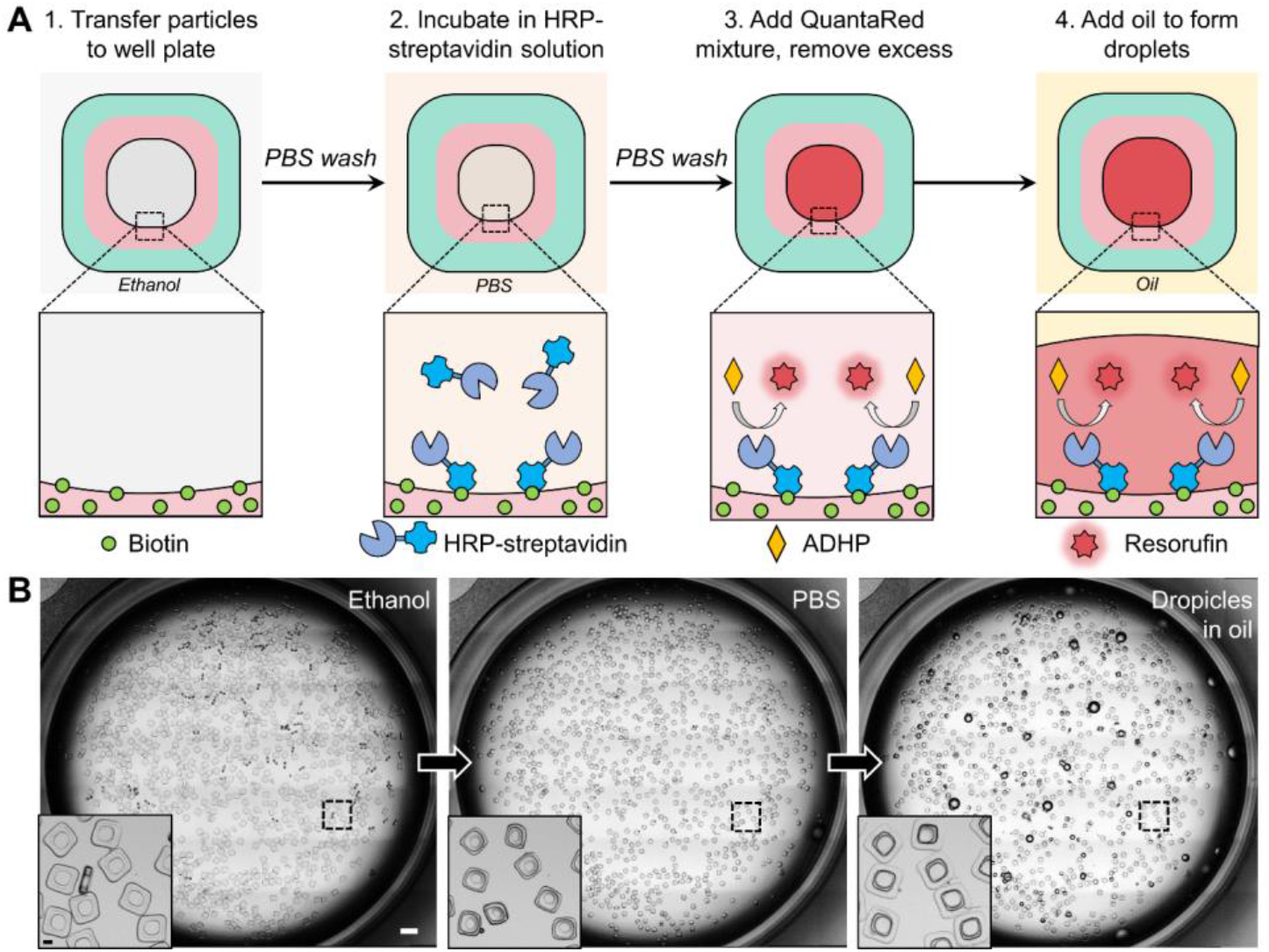
Amplified bioassay in dropicles. **(A)** Schematic of the assay workflow for biotin streptavidin affinity and the mechanism of amplification using horse radish peroxidase (HRP) turnover of the fluorogenic substrate ADHP to generate resorufin. **(B)** Microscopic images of a single well at different steps of the assay workflow. Images are captured when particles are in ethanol (step 1), PBS (step 2), and PSDS oil (step 4) where dropicles are formed. Zoomed in inset images of the same field-of-view highlight the particle morphology changes as the inner PEG and outer PPG layers swell or shrink in different solutions. Scale bar for the whole well is 1 mm, and the scale bar for the insets is 100 μm.

We observe selective amplification within dropicles and minimal cross talk of signals between particles functionalized with an affinity moiety (biotin) and those without. Plus-shaped particles without biotin in the PEGDA layer are used as a negative control population while H-shaped particles with biotin in the PEGDA layer are used as a positive population with high affinity to streptavidin-HRP (**Figure 4A**). Upon incubation together with a relatively high concentration of streptavidin-HRP (0.1 nM) the two populations are easily distinguished in both bright field, based on the structure of polymerized polymer, and fluorescence images (Figure 4B-C). The fluorescent signal from plus-shaped particles (negative group) increases at a much slower rate compared to that of H-shaped particles (positive group) (Figure 4D). The average intensity at 15, 35, and 60 min time points is 9.0, 12.1, and 12.3 fold higher for biotin-modified particles compared to non-modified particles, respectively (Figure 4E). The fluorescence intensity differences for these two mixed particle types over time is consistent with observations for the same particle types that were not mixed together, indicating minimal transport of the produced resorufin dye through the oil phase. Even at 48hrs, although some of the droplets are partially evaporated, the signals from these two particle types remain noticeably different (Figure S5). These results demonstrating limited cross talk support the potential for multiplexed detection using shape-coded particles.

**Figure 4.**
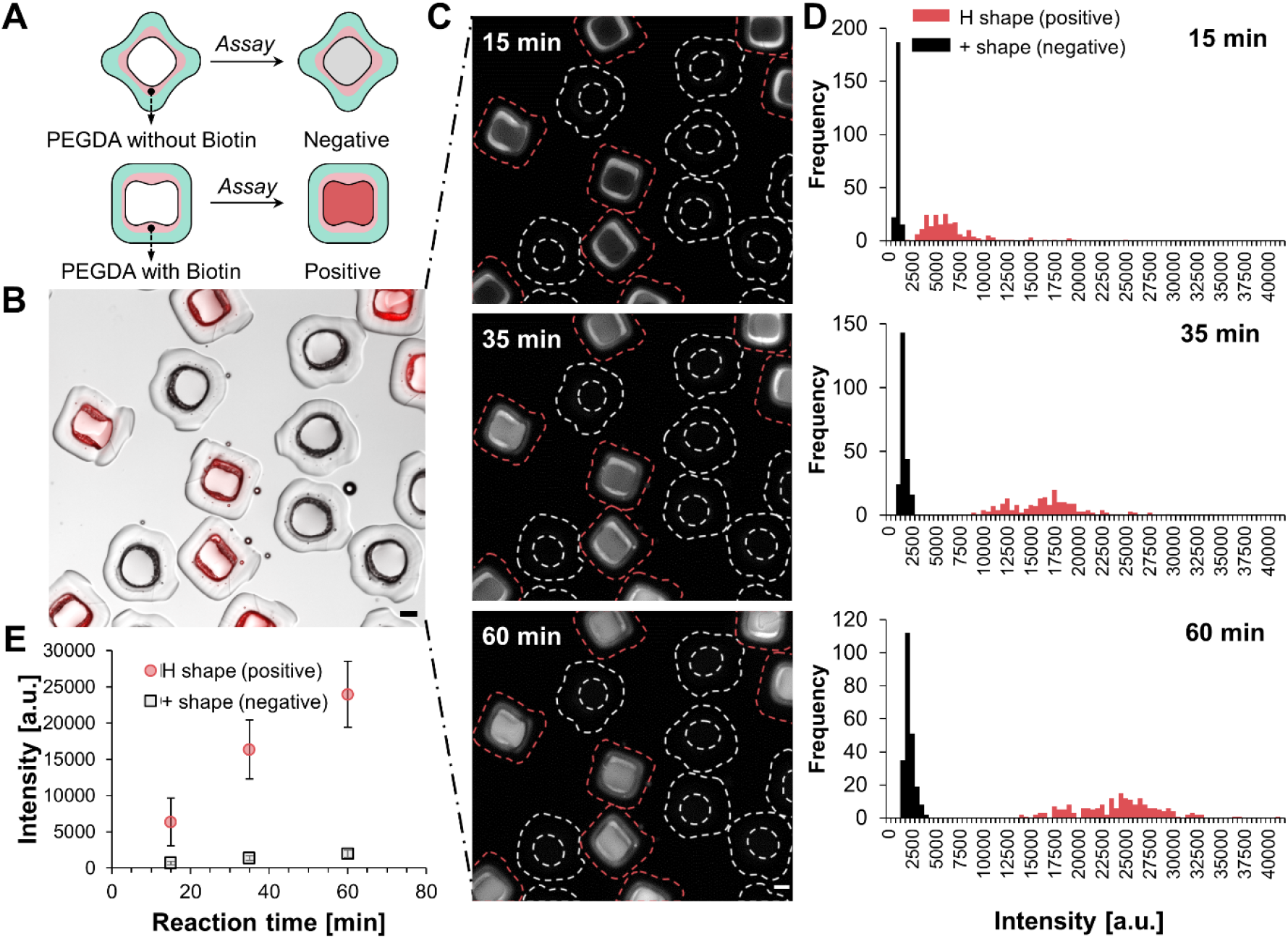
Duplex assay using shape-coded particles demonstrating minimal cross talk. **(A)** Schematic of the duplex assay showing the two particle populations: plus-shaped particles without biotin in the PEGDA layer are used as negative control population; H-shaped particles with biotin in the PEGDA layer are used as a positive population. **(B)** A merged image of bright field and fluorescent channels, after mixing plus- and H-shaped particles, incubating with 0.1 nM streptavidin-HRP solution, and initiating the HRP amplification reaction. There is contrast in the red fluorescent signal between plus-shaped particles and H-shaped particles. **(C)** Fluorescence images of the same field of view as in (B) at 15, 35 and 60 min after initiating the reaction. Red dotted lines in the images outline the PPG boundary of H-particles (positive) while white dotted lines outline the PPG boundary of plus-particle (negative). Scale bars in (B-C) represent 100μm. **(D)** Histograms showing the intensity distribution for a population of plus-shaped and H-shaped dropicles at 15, 35 and 60 min. **(E)** The mean of fluorescent intensity at these three timepoints 15, 35, and 60 min for H- and plus-shaped particles, showing the negative volume does not appear to have an appreciable increase in intensity over time even as the positive volume shows high levels of fluorescence intensity increase. The error bars represent standard deviation.

Using an amplified assay in dropicles we achieve a sub-picomolar level detection limit while maintaining a wide dynamic range. In addition, the solid substrate templating the droplet provides flexibility in adjusting the number of binding sites per particle. We report results from a concentration sweep of streptavidin-HRP for two biotinylated particle types. Particle Design 1 has an extruded height of 200 μm and a ~2-fold thicker PEGDA layer compared to particle Design 2 which has a height of 100 μm and thinner PEGDA layer (**Figure 5A**). Both particle designs achieved <1 pM detection limit with an at least 4-orders-of-magnitude dynamic range (Figure 5B-C). In comparison, Design 1 exhibits a detection limit of 100 fM, and linear range from 1 pM to 1 nM (3 orders of magnitude). Design 2 exhibits an improved detection limit of 10 fM, but a narrower linear range from 1 pM to 100 pM (2 orders of magnitude). The detection limit was experimentally determined as the lowest statistically differentiable concentration (with > 99.9% confidence level, i.e., *p* < 0.001 using student’s t-test) between the population of particles with droplets containing an analyte concentration versus droplets exposed to the same workflow but without analyte. Notably, amplified assays in dropicles from both designs outperformed direct binding of fluorescent streptavidin to the particles (Figure S6). Direct binding of streptavidin-Alexa Fluor^®^ 568, using the same particles as shown in Figure 5C, led to a weaker signal, requiring at least 1nM of streptavidin for detection, equivalent to a 5-orders-of-magnitude signal enhancement with the same number of binding sites per particle. This suggests that signal amplification is important to maximize the capabilities of particle-based assays. Furthermore, the amplified dropicle assay flow was adapted to a sandwich ELISA workflow for the detection of N-terminal propeptide B-type natriuretic peptide (NT-proBNP), a guideline recommended biomarker for cardiovascular disease (Figure 5D). The dropicle system achieved a detection limit of 10 pg/ml (i.e., 285 fM) and the highest sensitivity between 100 pg/ml and 1 ng/ml which corresponds well to the clinically relevant range around a few hundred pg/ml.

**Figure 5.**
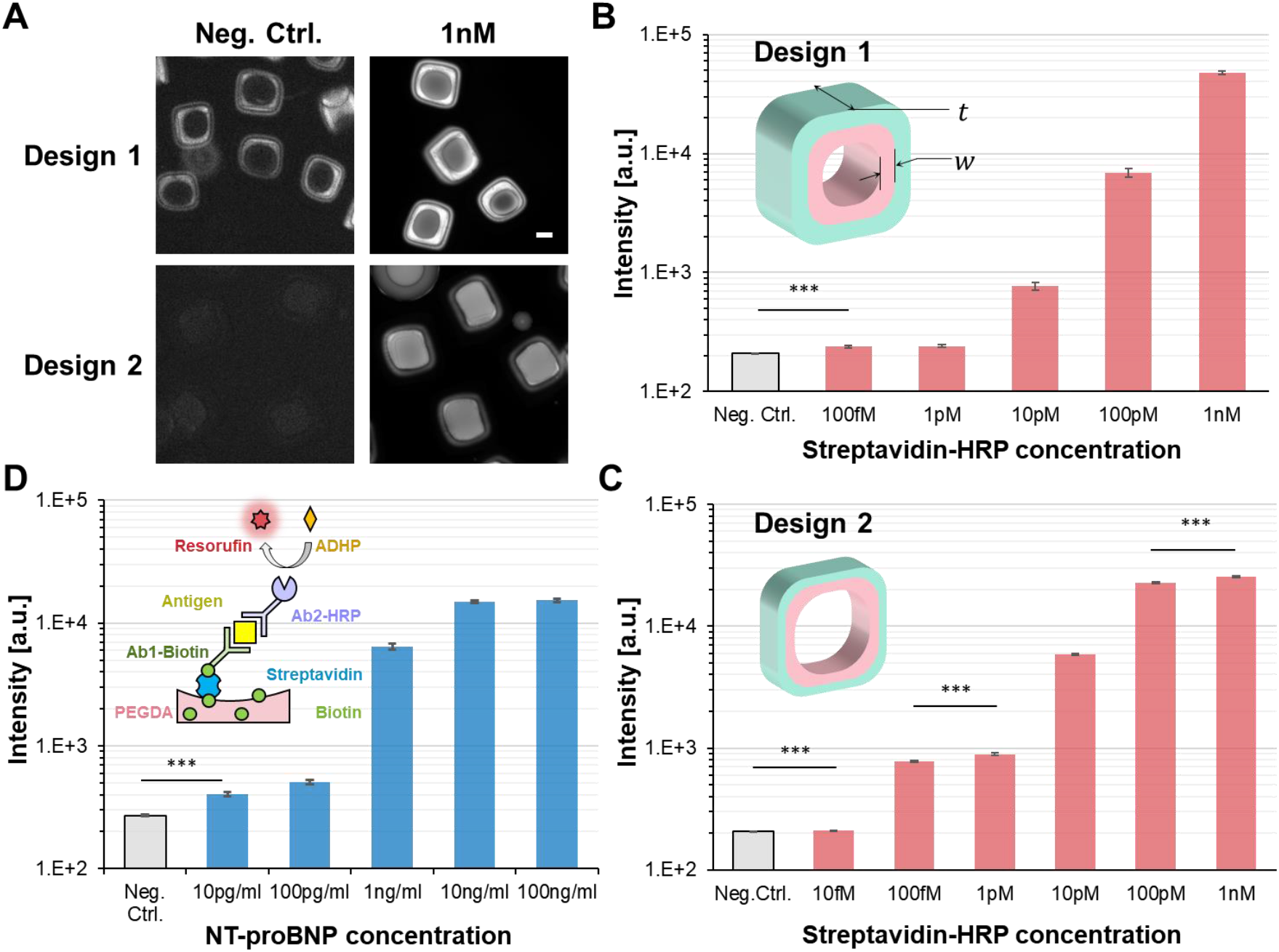
Amplified assay performance in dropicles. **(A)** Microscopic images of QuantaRed assay results using two O-shaped particle designs for two concentrations of streptavidin-HRP, i.e., negative control and 1 nM. The look-up table was adjusted to be the same for each condition for visibility. The scale bar represents 100 μm. **(B-C)** Mean amplified intensity across a population of dropicles as a function of concentration of streptavidin-HRP using (B) particle design 1 (N=11-38) and (C) particle design 2 (N=307-382), each tuned by varying the thickness (*t*) and width (*w*) of the internal PEG layer. **(D)** Mean amplified fluorescent intensity across a population of dropicles as a function of concentration of NT-proBNP. Data reported in (A-D) were obtained after 45 min of reaction. Error bars in (B-D) represent standard error. *** represents *p*<*0.001*.

## 3. Discussion

### 3.1. Instrument-Free Formation of Uniform Compartments

The dropicle system presented here provides a platform to create uniform nanoliter-scale compartments with existing laboratory equipment and processes. Analyte-specific batches of different shape-coded particles can be manufactured in bulk and easily distributed to life science researchers interested in performing compartmentalized bioassays. We estimate that the material cost to produce 15,000 particles, more than enough for an assay, to be ~$1 (Supporting Information). There is no need for microfluidics experience or specialized equipment because droplet formation requires only simple pipetting steps typical of a standard ELISA assay workflow. Two additional steps specific to a dropicle workflow are: 1) An initial transfer and media exchange step such that amphiphilic particles stored in ethanol are resuspended in an aqueous solution. 2) A final step following addition of a readout solution, such as QuantaRed, in which excess solution is removed immediately from the well plate and oil is added on top of the particles to seal the droplets formed. Both of these steps are simply achieved using typical pipetting techniques within 1-2 min, therefore no additional training or instruments are needed for novice users to implement sophisticated nanoliter-scale compartmentalized reactions. For more experienced researchers interested in fabricating the reported particles or customizing them, the 3D printed device designs we use, which we have made freely available (Supporting Information), can be outsourced to a number of vendors and received within a few days. For customized particle designs, users can tune the co-axial channel cross-sections and the photomask to change the shape and thickness of the particles, respectively.

### 3.2. Improved Detection Accuracy Through Swarm Sensing

Hundreds of dropicles can be read in a single well, enabling swarm sensing.^[42]^ Biosensing accuracy suffers from low signal above background at low analyte levels and random variations in sensor performance at higher analyte levels which limit quantitation. Conventional detection schemes, such as ELISA, overcome low analyte level challenges through enzymatic signal amplification, however these assays typically measure the bulk signal from a single or few reactions, leading to compromised detection sensitivity and accuracy. When reading out reactions in dropicles, signal intensity is affected by a number of random factors including variations in particle manufacturing, variations in droplet volume, non-specific binding, and measurement noise introduced by the excitation and readout system. Accumulated random errors are significantly reduced by compiling a histogram or summary statistics of the independent signals from a large number of particles in the well (i.e., larger swarm size, Figure S7A). Accumulating these data results in a statistically robust determination of the ground truth signal, which can lead to a more robust and accurate quantification of concentration. For example, when sample size was increased from n=3 dropicles to n=300, the standard error, which is a measure of the accuracy of the sample mean compared to the population mean, was significantly improved (i.e., decreased ~7-16 fold for various concentrations with an average of ~12 fold, Figure S7B). Using information about expectations for the distribution of results for a given ground truth can also yield enhanced prediction capabilities ^[45]^. Depending on the application, the sample size (i.e., number of particles per well) could be further scaled up if higher sensing accuracy and detection fidelity are required.

### 3.3. Multiplexing Immunoassays in Dropicles

Multiplexed detection using microfluidic-based approaches, such as microbeads encapsulated in droplets, are limited in scale by multi-Poisson statistics. The loading efficiency of droplets with a desired combination of barcoded solid phase particles (i.e. only a single type per droplet) is expected to decrease drastically as the number of targets in the multiplexed panel increases. This inevitably escalates the imaging acquisition burden and prolongs readout time with the need to isolate potential combinations of barcodes for data analysis.^[46]^ A potential solution is to further enhance the throughput of the droplet generator and reader^[4]^ to shorten the time needed to accumulate a sufficient number of droplets containing only one particle of each single type. Our dropicle system provides an alternative approach to overcome this obstacle, as the single solid phase is designed into the amphiphilic particles during manufacturing, and therefore not subject to Poisson loading statistics. Moreover, the flexibility of particle design using customizable co-axial flow allows for the fabrication of a vast collection of particles with different outer/inner layer shapes and thicknesses. By using shape-coded particles, we only require signal readout in a single fluorescence channel together with bright field imaging for multiplex detection, alleviating the need for more complex multi-channel fluorescence readers and the challenges with compensation between fluorescence channels. Therefore, the shape-coded dropicle system is particularly beneficial for cost-effective assays that are compatible with similarly cost-effective readers based on consumer electronic devices,^[47,48]^ which paves the way for multiplexed detection at point-of-care settings or other limited resource settings.

Furthermore, modifying the dimensions (e.g. *t* and *w* denoted in Figure 5B) and potentially chemistry of the PEGDA layer^[21,29,38]^ allows for tuning of the detection limit and dynamic range of an assay for the optimal detection of target analytes in each dropicle. For disease diagnosis, multiplexed *in vitro* assays are often limited by the capability to detect analytes spanning a wide range of clinical cutoffs. For instance, among cardiac biomarkers, the clinical cutoffs range from pg/ml (e.g., cardiac troponins), sub-ng/ml (e.g., natriuretic peptides), ng/ml (e.g., creatine kinase-MB), to μg/ml (high-sensitivity c-reactive protein).^[49]^ To tackle this issue, commercial assays and research platforms focus on the development of either a single assay with a wide dynamic range^[50]^ with trade-offs in quantitative accuracy, or a multi-modal sensing approaches to cover different concentration ranges.^[51,52]^ Tuning the assay characteristics within each type of barcoded dropicle promises a simpler and more practical approach to achieve the simultaneous detection of multiple markers. In addition, a collection of different shape-coded particles tuned for quantitative accuracy in a particular concentration range could also be utilized to expand the dynamic range for detection of a single marker, minimizing the need for sample enrichment or dilution in a clinical workflow.

### 3.4. Lab on a Particle Systems

By enabling the simple formation of uniform and stable nanoliter compartments in a biocompatible oil phase without cross talk, dropicles can serve as a platform for a variety of molecular and cellular assays leveraging standard laboratory equipment. The enzymatic affinity assay we demonstrate is a foundational component of many ELISA workflows, indicating dropicles should be well-suited for biomarker detection using immobilized antibodies, aptamers, or other affinity elements for *in vitro* diagnostics ultimately approaching single molecules.^[53]^ Our system maintains the advantages of amplification that is achievable with bulk assays, while also providing unique benefits of barcoded particle-based assays and smaller volume analysis. Furthermore, it enables rare or low volume sample multiplexed analysis all in one well using enzymatically amplified assays, which is not possible to achieve without the compartmentalization capability. Importantly, analysis of hundreds to thousands of reactions in a single well could lead to the ability to perform more tests with limited sample volume, reduce reagent consumption, and overall assay time, compared to conducting assays in large volume wells. These benefits are particular attractive for clinical trials, drug screening, and diagnostics, where sample volumes can be small or precious.

Digital nucleic acid detection using compartmentalized amplification of single nucleic acids with PCR or Loop-mediated isothermal amplification (LAMP)^[54]^ should also be compatible with dropicle systems given the similar volumes to current systems and biocompatibility of the PEG layer and surrounding oil. In addition, by modifying the PEGDA layer with cell adhesive-moieties such as biotinylated collagen,^[38]^ our amphiphilic particles could serve as cell carriers, enabling single cell culture and analysis, including the accumulation of secretions^[55]^ or RNA in the small volume of a dropicle for secretion or gene expression analysis.^[56]^ Building off this work, we envision such dropicle systems can serve as minimally-instrumented and accessible “lab on a particle” platforms for analysis of molecules and cells at the ultimate limits of biology.

## 4. Experimental Section

### 4.1. Particle Manufacturing Setup

The particle manufacturing system comprises a fluidic system with a 3D printed microfluidic device connected to syringe pumps to drive the pre-polymer co-flow and a pinch valve to stop flow, and an optical system in which the co-flow is exposed to patterned UV light through a mask. The 3D printed microfluidic device connected to a glass capillary is fixed on top of a custom-built stage. The inlet ports are connected (PEEK Union Assembly P-702, OD 1.58 mm, IDEX, IL, USA) to four syringes (20 ml Plastic Syringe with Luer-Lok Tip, BD, NJ, USA) mounted on two separate syringe pumps (PHD 2000, Harvard Apparatus, MA, USA) using PTFE tubing and Luer stubs. PTFE tubing from the outlet Luer stub is passed through a pinch valve (2-Way Pinch Valve, SCH284A003, ASCO, NJ, USA) into a collection vessel (conical tubes, Corning, NY, USA). A photomask (Chrome Film Mask, CAD/Art Services, OR, USA) is taped on top of the glass capillary to provide a controlled UV exposure. A UV source (OmniCure S2000, Excelitas Technologies, MA, USA) exposes the capillary’s region of interest under the photomask through a light guide with a collimator (Adjustable Spot Collimating Adaptor, Excelitas Technologies, MA, USA) and a light shutter (Lambda SC, Smart Shutter control system, Sutter Instrument, CA, USA) attached at its end. The syringe pumps, valve and shutter are automated and controlled using a graphical user interface developed in LabVIEW (National Instruments, TX, USA).

### 4.2. Particle Fabrication

Four different streams of density matched solutions are pumped through the 3D printed device inlets 1-4 at flow rates *Q*_*1*_ to *Q*_*4*_, respectively with the syringe pumps, yielding an inlet pressure of *P*_*in*_ (Figure 1A). The net flow rate results in an average velocity of *U*_*avg*_ within the square section of the glass capillary. The pinch valve at the outlet is used to stop the flow entirely (*U*_*avg*_ = 0) such that the outlet pressure of *P*_*out*_ (= *P*_*in*_), whereas the syringe pumps are simultaneously turned off to cut the flow through the inlets. The shutter is opened after a short delay time (*τ*_*d*_) to ensure that the flow has fully stopped, whereas the UV light source with intensity *I*_*UV*_ exposes the flow streams through a photomask for an even smaller exposure time (*τ*_*exp*_) (Figure 1C). Co-axial flow streams are exposed to rectangular-shaped UV patterns defined by an array of 20 or 30 slit features (width 100 or 200 μm × length 1000 μm) in the photo mask. The width of the UV beam defines the thickness of the polymerized region and the particle, whereas the 1000 μm breadth of the UV beam is enough to cover the whole cross-section of the rectangular glass capillary. The extended breadth of the pattern (1000 μm) also allows for easy alignment with the glass capillary (700 μm outer dimension). The flow streams originating from channels 2 and 3 are polymerized in the form of a ring-shaped amphiphilic particle with hydrophobic outer layer made of PPGDA and hydrophilic inner layer made of PEGDA (Figure 1D). At this point, the pinch valve is opened (*P*_*out*_ = 0) and the syringe pumps are started again to reach an average flow velocity of *U*_*avg*_ within a flow stabilization time (*τ*_*s*_). The whole cyclic process of flow stoppage, UV exposure and flow stabilization is further described by plotting *P*_*in*_, *P*_*out*_, *I*_*UV*_ and *U*_*avg*_ against time and indicating *τ*_*d*_, *τ*_*exp*_, and *τ*_*s*_ in Figure 1B. One fabrication cycle is completed within ~5s for most experimental conditions. An automated experimental setup integrated through a LabVIEW GUI allows for rapid and on-demand control of the parameters (*τ*_*d*_, *τ*_*exp*_, *τ*_*s*_ and *U*_*avg*_). The average flow velocity is controlled by adjusting the flow rates for the four inlets *Q*_*1-4*_.

Production conditions for example O-shaped particles shown in Figure 1F are provided. Particles were fabricated by pumping the polymer precursors through the four separate inlets of the 3D printed device at flow rates of 0.25 ml/min each, where *τ*_*d*_ = 1.25 s, *τ*_*exp*_ = 0.3 s, and *τ*_*s*_ = 4 s. A net flow rate of 1ml/min results in *U*_*avg*_ = 66.7 mm/s, Reynolds number *Re* ≅ 1.4 and Peclet number *Pe* = 3.3 × 10^5^ for a glass capillary with hydraulic diameter of *d*_*h*_ = 0.5 mm. For a UV exposure length of ~10 mm, it takes *t* ≅ 0.15 s for a fluid particle to travel such a distance. Therefore, the diffusion length 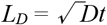 for the PI molecules is calculated as 3.9 μm with a diffusion coefficient *D* = 10^−10^ m^2^/s. Diffusive blurring at this length scale results in a minor variation (<1%) in the PI concentration profile compared to the microchannel width (~0.5 mm) and length (~10 mm). Conditions for manufacture of other particles used are described in the Supporting Information.

### 4.3. Dropicle Formation

Dropicles are formed using a simple workflow based on pipetting and washing (Figure 2A). Particles initially suspended (and stored) in ethanol (step 1) are transferred to a hydrophobic well plate where the medium is exchanged to PBS after three washes (step 2). Introduction of an aqueous solution results in swelling of the inner PEG layer thus reducing the inner cavity size to some extent. To characterize the uniformity of droplet volume in dropicles, 20 μg/ml biotin-4-fluorescein (AnaSpec, CA, USA) dissolved in PBS is added (step 3). Once the aqueous fluorophore solution is fully dispersed around and inside the particle cavity by wetting the hydrophilic inner surface, excess liquid is removed (step 4) while an aqueous phase remains trapped within particle cavities. Poly(dimethylsiloxane-co-diphenylsiloxane) (PSDS, Sigma-Aldrich, MO, USA) oil is added on top of the particles to complete the compartmentalization of the aqueous phase within the hydrated particles by pushing any remnant aqueous phase outside of the particles away from them (step 5). After the oil is added, the particles gradually recover back to their original shape as the PSDS swells the outer PPG layer in a similar manner as the original ethanol storage solution.

### 4.4. Amplified Affinity Assay with QuantaRed

Particles suspended in ethanol are first added to a 12 well plate. Once particles settle in the well excess ethanol is removed, followed by three washes with PBS with 0.5% w/v Pluronic (PBSP). For each washing step, 500μL of washing buffer is added to the center of the well, and subsequently removed from the corner of the well. This process is repeated three times to complete each wash cycle. Then, 300 μl of streptavidin-HRP solution at varying concentrations (Thermo Fisher Scientific, MA, USA) is added and incubated for a given time period, followed by three additional washes with PBSP. Next, 500 μl QuantaRed solution (Thermo Fisher Scientific, MA, USA) is mixed following the instructions of the vendor (at a 50:50:1 ratio of enhancer solution, stable peroxide, and ADHP concentrate, respectively) and added to the well to wet the particles with excess removed immediately. Lastly, 500 μl oil (PSDS) was added to form isolated dropicles. Next, fluorescence and bright field images of the dropicles in oil are obtained at desired time points using a fluorescence microscope. The excitation and emission maxima of the fluorescent product are at ~570 and 585 nm, respectively. Imaging of the whole well allows the simultaneous monitoring of a few hundreds of reactions in compartmentalized dropicles. The same assay protocol was used for assay performance characterization using two types of O-shaped particles. In the negative control group, particles were incubated with PBS only, all the other steps were kept the same as positive groups as described in Figure 3. To determine the fluorescence intensity of the amplified assay, region-of-interests (ROI) are defined within each droplet held within the dropicle (excluding the inner PEG layer), and the average intensity was extracted using ImageJ to represent the amplified signal of each dropicle. The very small fraction of particles that are settled on their sides have distinctive shape differences, which allows them to be differentiated from particles facing upwards and excluded from analysis using parameters such as circularity and area.

### 4.5. Duplex Experiment

Plus-shaped particles without biotin in the PEG layer are used as a negative control population; H-shaped particles with biotin in the PEG layer are used as a positive population. These two types of particles have different shapes of the outer PPG layer and inner PEG layer, and can therefore be easily distinguished by shape in both bright field and fluorescence channels. Both types of particles are mixed at a 1:1 ratio in ethanol in an Eppendorf tube and subsequently transferred to a well in a 12 well plate. Particles are then washed with PBSP three times, and incubated with 0.1 nM streptavidin-HRP solution for 30 min. Then, the same QuantaRed assay protocol is performed as described above, where particles are washed again, and droplets are subsequently formed encapsulating the QuantaRed mixture.

### 4.6. NT-proBNP Detection Using Amphiphilic Particles

We developed a sandwich ELISA for NT-proBNP detection using commercial monoclonal antibodies. A monoclonal capture antibody (15C4cc, HyTest, Finland) was conjugated with biotin, and a monoclonal detector antibody (13G12cc, HyTest, Finland) was conjugated with HRP, respectively, using commercial conjugation kits (Lightning-Link, Expedeon, United Kingdom) following vendor instructions. Particles with biotin were transferred to a well plate in ethanol and washed as described above. To immobilize capture probes onto the particles, 10 μg/ml streptavidin (Thermo Fisher Scientific), MA, USA was added to the well and incubated for 30 min, followed by three washes. Then, particles were incubated with 10 μg/ml biotin conjugated capture antibody for 1 hr to complete the immobilization step, where unbound antibodies were removed through washing steps. Next, blocking buffer (Thermo Fisher Scientific, MA, USA) was added to the well containing particles and incubated for 1 hr followed by washing. To test the detection of NT-proBNP, particles were incubated with varying concentrations of human recombinant NT-proBNP (HyTest, Finland) for 1 hr with subsequent washing, followed by incubation with 0.5 μg/ml HRP conjugated detector antibody for 1 hr, and a final washing step. Lastly, the QuantaRed assay solution was added for amplification, and droplets were formed in PSDS oil as described above. In the negative control group, particles were incubated with PBS only instead of NT-proBNP, but all the other assay steps, including immobilization, blocking and detection etc., were kept the same as the positive groups. Fluorescent intensity was measured 45 minutes after adding QuantaRed solution.

## Acknowledgements

This project is supported by the NSF-Engineering Research Center for Precise Advanced Technologies and Health Systems for Underserved Populations (PATHS-UP) - Award Number 1648451. We also acknowledge the Simons Foundation Math+X Investigator Award #510776.

## Supporting Information

### 1. Laminar Flow and Diffusion Simulation

The laminar fluid flow inside the 3D printed devices and diffusion of the PI across the streamlines are simulated by coupling “Laminar Flow” and “Transport of Diluted Species” modules in COMSOL Multiphysics, MA, USA (Figure 2B). A single phase fluid (density 987 kg/m^3^ and viscosity 30 mPa.s) flow is solved for as a stationary case. The viscosity of the fluid is assumed to be constant throughout the fluid domain, whereas a diffusion coefficient value of 10^−10^ m^2^/s for the PI is used. The inlet boundary conditions are set as laminar flow rates of 0.25 ml/min for all the inlets, whereas the outlet boundary condition is set at atmospheric pressure (0 Pa). Symmetric boundary conditions are used wherever applicable. The solution time for this problem with a 3D fluid domain is approximately 2 hrs on a computer with 8GB RAM and 2.33 GHz Core 2Duo processor. The experimental viscosities of the density matched PEGDA in ethanol and PPGDA in ethanol solutions are measured as 7.55 mPa·s and 38.3 mPa·s, respectively. However, solving a two-phase flow, with different viscosities for each fluid, by coupling the “Laminar Two-Phase Flow, Phase Field” and “Transport of Diluted Species” modules is computationally expensive for a 3D geometry. Therefore, we instead solve a 2D axisymmetric case including the viscosity differences in the fluids. The results of this simulation showed a reasonable agreement in diffusion characteristics of the PI across the microchannel width. The solution time for the axisymmetric problem is more than 20 hrs. For brevity, the results for the axisymmetric case are not reported here.

The shapes of the fabricated particles match well with simulation results predicting the PI concentration distribution across the channel width and the location of the streamlines corresponding to each polymer precursor (Figure S1B, Figure S2). Diffusive blurring of the PI distribution leads to smoothening of sharper gradients of PI and better reflects the actual particle morphologies (Figure S1B). However, these blurred shaped particles do not affect the performance of the assay. Moreover, to overcome the effect of diffusive blurring and achieve better particle definition, the outer and inner sheath flow are replaced with inert non-crosslinkable precursors mixed with similar concentrations of PI. Therefore, the shape of the particles is less effected by the diffusion and is more accurately defined by the fluid streamlines as indicated in Figure S2.

### 2. Droplet Characterization

It is estimated that the O, H, Plus, and U shapes held approximately 2.8, 3.5, 4.1, and 2.9 nanoliters, respectively, largely due to differences in their internal PEG layer geometry (Figure S1C, Figure S1E). We also estimate the uniformity in drop size by calculating the variation in intensity of an encapsulated fluorophore. We calculated the drop volume for seven different O shaped particles with varying diameters and thicknesses, and obtained a CV of 9.3% on average. Assuming a spherical shape for a droplet this would only correspond to an approximately 3% CV in droplet diameter.

### 3. 3D Device Design

A 3D microfluidic device, having four inlet ports and one outlet port, is designed in AutoCAD (AutoDesk, CA, USA). Different devices are designed separately with various cross-sections of the channels for similar-shaped particles fabrication (Figure 2, Figure S2). The inlet ports lead to four stacked microfluidic channels (635 μm) separated by a thin wall (400 μm). All the channels are sequentially merged together within a 3-4.5mm long tapered region close to the outlet port, where the outer most dimension is reduced from ~9 mm to ~0.7 mm. First the inner most channel is joined with the adjacent channel by removing the first wall in between them. As the cross-sectional dimension of the tapered region reduces further, the next channel is joined with the first two by removing the second wall in between them. Finally, when the tapered region converges close to the outlet port dimension, the fourth channel is also merged together with the first three channels. At the exit of the 3D printed part, the outlet cross-section is designed as a tapered square shape (0.7 mm × 0.7 mm to 1 mm × 1 mm) so that a square glass capillary (ID 0.5 mm, OD 0.7 mm) could be easily fit and align with the co-axial channels. The inlet ports have a diameter of 1.58 mm to tightly connect with the Polytetrafluoroethylene (PTFE) tubing with OD 1.58 mm. The microfluidic device is 3D printed using a photopolymer (WaterShed XC 11122, ProtoLabs, MN, USA) and a high-resolution (50 μm layers) stereolithography technique (ProtoLabs, MN, USA). The square glass capillary (8250, 50 mm, VitroCom, NJ, USA) and the PTFE tubing (Kimble, OD 1.58 mm and ID 0.78 mm, DWK Life Sciences, Germany) are glued (Devcon 5-Minute Epoxy 20845, ITW Consumer, CT, USA) to the 3D printed part (~11 mm × ~15 mm × ~21 mm) and placed on top of a stack of supporting glass slides. A Luer stub blunt needle (LS21, Instech Laboratories, PA, USA) is also glued at the end of the glass capillary.

### 4. Materials for Particle Fabrication

For the hydrophilic and hydrophobic layers of the multi-material 3D particles, poly(ethylene glycol) diacrylate (PEGDA, *M*_*w*_ ≈ 575; 437441, Sigma-Aldrich, MO, USA) and poly(propylene glycol) diacrylate (PPGDA, *M*_*w*_ ≈ 800; 455024, Sigma-Aldrich, MO, USA) are chosen to be the polymer precursors, respectively. The densities of the PPGDA (1.01 g/cm^3^) and PEGDA (1.12 g/cm^3^) solutions are matched (0.987 g/cm^3^) by adding 10% and 40% ethanol (0.789 g/cm^3^) in the mixtures, respectively. The photoinitiator (PI) concentration for channel 2 and 3 is maintained at 5% of the total volume of the PPGDA (90%) in ethanol (10%) and PEGDA (60%) in ethanol (40%) mixtures, respectively. For flow configuration 1, the photoinitiator (2-hydroxy-2-methylpropiophenone, Darocur 1173, 405655, Sigma-Aldrich, MO, USA) is added only to the two precursors that are to be polymerized upon UV exposure, when the outer and inner sheath flows are PPGDA and PEGDA, respectively. However, for flow configuration 2, to reduce the effect of PI diffusion on particles blurring, the outer and inner sheath flows are replaced with PPG (*M*_*w*_ ≈ 400; 81350437441, Sigma-Aldrich, MO, USA) mixed with PI and PEG (*M*_*w*_ ≈ 200; P3015, Sigma-Aldrich, MO, USA) mixed with PI, respectively. For the biotin to streptavidin-HRP binding assays reported, the inner PEGDA layer is also biotinylated to enable binding of streptavidin. Before the particle fabrication, 0.25 ml of acrylate-PEG-biotin (APB, PG2-ARBN-5k, NANOCS, NY, USA) dissolved in DMSO (100 mg in 1.66 ml) is mixed with 20 ml of PEGDA and ethanol solution. The biotin is grafted within the PEGDA layer during photo-crosslinking.

For a final concentration of 0.75 mg of APB per ml of PEGDA solution, we can estimate 9.033×10^16^ APB molecules/ml or 9.033×10^16^ APB molecules/m^3^. A 5 kDa APB molecule has an approximate 1.1 nm radius of gyration or 2.2 nm diameter. An APB molecule would be available for binding if it is present within a thickness of < 2.2 nm from the particle’s inner cavity surface exposed to the aqueous phase. For an O-shaped amphiphilic particle with a cavity diameter of 200 μm, and thickness of 100 μm, we estimate the number of APB molecules present within a 2 nm (< 2.2 nm molecular diameter) PEGDA width to be ~1.1 ×10^11^ per particle.

### 5. Experimental Conditions for Particle Fabrication

The experimental conditions for the manufacture of the particles using flow configuration 1 described in Figure S1C are as follows: for O-shaped particles, *τ*_*exp*_ = 0.3 s, *τ*_*d*_ = 1.25 s, *τ*_*s*_ = 4 s, and total flow rate (*Q*_*t*_ = *Q*_*1*_ + *Q*_*2*_ + *Q*_*3*_ + *Q*_*4*_) = 1 ml/min, *Pe* = 3.3 × 10^5^; for plus-shaped particles, *τ*_*exp*_ = 0.3 s, *τ*_*d*_ = 1.75 s, *τ*_*s*_ = 2 s, and *Q*_*t*_ = 2 ml/min, Pe = 6.6 × 10^5^; for H- and U-shaped particles, *τ*_*exp*_ = 0.3 s, *τ*_*d*_ = 2.25 s, *τ*_*s*_ = 4 s, and *Q*_*t*_ = 1 ml/min, Pe = 3.3 × 10^5^. The average inner dimensions of the particles for the above experimental conditions are measured as 187.6 ± 6.7 μm, 209.9 ± 5.7 μm, 228.7 ± 2.6 μm, and 194.4 ± 6.1 μm for O-, H-, Plus-, and U-shaped particles, respectively. The flow rates ratios are kept constant for these experiments (*Q*_*1*_ = *Q*_*2*_ = *Q*_*3*_ = Q_*4*_ = 1). It is worth noting that the flow stabilization time could be reduced to half, *τ*_s_ = 2 s, if the flow rate is doubled (*Q*_*t*_ = 2 ml/min). The experimental conditions corresponding to the four different flow rate ratios of 1 to 4 in Figure S1D are as follows: *τ*_*exp(1-4)*_ = 0.3 s, *τ*_*d(1-4)*_ = 1, 1.25, 1.5, 1.75 s; *τ*_*s(1-4)*_ = 4 s, and *Q*_*t(1-4)*_ = 1, 1.5, 2, 2.5 ml/min. The experimental conditions for fabrication of particles used in assay experiments (Figure 5) are as follows: (design 1) PI concentration of 5%, *τ*_*exp*_ = 0.3 s, *τ*_*d*_ = 1 s, *τ*_*s*_ = 4 s, and *Q*_*t*_ = 1 ml/min (*Q*_*1,2*_ : *Q*_*3,4*_ = 1), particle’s thickness defined by the mask width *t* = 200 μm; (design 2) PI concentration of 5%, *τ*_*exp*_ = 0.3 s, *τ*_*d*_ = 1.75 s, *τ*_*s*_ = 2 s, and *Q*_*t*_ = 2 ml/min (*Q*_*1,2*_ : *Q*_*3,4*_ = 1), *t* = 100 μm. The experimental conditions for the manufacture of the particles with flow configuration 2 described in Figure 2 are as follows: *τ*_*exp*_ = 0.5 s, *τ*_*d*_ = 1.5 s, *τ*_*s*_ = 4 s, and *Q*_*t*_ = 1.5 ml/min, *Pe* = 4.95 × 10^5^.

### 6. Multi-material Particle Characterization

The particles collected directly from the device after a complete fabrication cycle are initially suspended in a mixture of uncured PEGDA and PPGDA in ethanol. Additional ethanol is added to reduce the overall density of the solution so much so that the particles becomes heavier than the media and settle on the bottom of the conical sample collection tube. After removing the supernatant, the particles are washed with pure ethanol three times with >100× volume and are stored in ethanol for later experiments. For PEGDA layer visualization, the particles are transferred from ethanol to phosphate-buffered saline (PBS) after three washing steps. The particles are incubated with resorufin dissolved in PBS buffer (100 μM solution) for ~10min. After washing the excess resorufin away with PBS (×3), resorufin partitioned into the PEGDA layer only is observed in the TRITC channel of the fluorescence microscope images. For characterization of particle dimensions, the fabricated particles suspended in ethanol are dispersed in a 12-well plate (Falcon untreated cell culture plates, Corning, NY, USA), where images are captured using a microscope and image analysis is performed using ImageJ (NIH, MD, USA). By adjusting the threshold, the inner and outer boundaries of each particle are identified, and the areas encompassed by these boundaries are measured separately. Correlating the measured area to the area of a circle, average diameters are deduced that represent the inner and outer dimension of the particle. In some case, a band pass filter is applied to the images before applying the threshold that helps in clearly identifying the boundaries of the particles.

### 7. Particle Retention Rate

For 300-600 seeded particles in a 12 well plate more than 98% of particles can be retained after the washing steps (data examined for five different experiments). To calculate the retention rate, the number of seeded particles were counted in ethanol and later in PBS, after all the washing steps were performed. High retention rates reflect the density differences between the multi-material particles and the surrounding media, and the exchange of media using gentle pipetting techniques without excessive agitation. When the particles are initially transferred to the well plate in ethanol, they naturally settle on the surface due to the slightly higher material densities (1.12 g/ml for PEGDA and 1.01 g/ml for PPGDA) compared to ethanol (0.789 g/ml). In the later assay steps such as biomolecule incubation and washing steps, conducted in aqueous solution (typically PBS, 1.005 g/ml), these particles with hydrophobic outer layers prefer the hydrophobic surface of the well plate in PBS, so they are not lost during the washing steps. Moreover, during the solution transfer steps, the solutions were gently added to the center of the well, and excess solutions were removed from the corner of the well. As the fluid is sucked from the corner side of the well, the induced shallow boundary layer flow profile does not exert a significant drag force to disrupt the already settled particles from their positions. For a higher seeding density, we observe 94.3% retention rate when the particles are transferred from ethanol to PBS and washed three times as shown in Figure 3 (1048 particles in ethanol, 989 particles in PBS), where the whole well image of hundreds of particles is monitored at different steps. In the inset images, the particles retained similar locations and orientations throughout the process. Finally, when the oil (1.05 g/ml) is added for encapsulation, it is absorbed by the outer hydrophobic polymer layer of the particle as well as priming the hydrophobic well surface. However, the particles stay settled at the bottom of the well, presumably because of the density differences as described above and the viscous nature of the oil that dampens any significant movements of the particles after the droplets are formed. These particle handling and medium exchange steps are very similar to the steps for conducting cell-based assays in a well plate, where excessive agitation is in general avoided to minimize unnecessary cell loss. On the other hand, we did observe that if agitation of the well plate is applied after droplet formation, the particles with droplets could be easily retrieved and transferred to another well or tubes as desired, which adds to the flexibility of downstream analysis.

### 8. Costs and Scalability

The pure polymer precursors, PPG, PPGDA, PEGDA, and PEG cost ~$78, ~$185, ~$180, and ~$25 per liter each. The density matched mixtures in ethanol (~$5/L) cost ~$71, ~$167, ~$110, and ~$17 per liter, respectively, averaging at ~$91/L. An addition of 5% PI ($1/mL) in each of the precursors cost ~$141/L on average. For an overall flow rate of 1 mL/min (i.e. 0.25 mL/min for each stream) and a continuous 4s of pumping after each UV exposure to create ~30 particles per cycle, it takes ~2.2 *μ*L of reagents per particle fabricated. A liter of reagents can be easily transformed into ~2.2 × 10^6^ particles. Therefore, more than 15,000 particles can be easily fabricated with ~$1 worth of reagents. Particles are manufactured in a continuous and automated manner which is scalable to large batches. The throughput of particle production is less important than the throughput of drop generation for microfluidic droplet assays because for microfluidic assays the total assay time includes the time to produce the assay droplets. Production time for particles occurs separately and is decoupled from the time to produce droplets. Dropicle formation occurs rapidly in less than 1 minute for an entire well filled with particles.

### 9. Direct Binding of Streptavidin to Particles

To evaluate signal from direct binding, particles (design 2) are transferred to a well plate and washed in the same manner as above followed by incubation with streptavidin-Alexa Fluor^®^ 568 (Thermo Fisher Scientific, MA, USA) at varying concentrations for 30 min, which is the same duration as for the amplification reaction, then three washes in PBSP. Next, dropicles were formed by adding oil. In the negative control group, particles were incubated with PBS only and all the other steps were kept the same as the positive groups. Fluorescence (TRITC) images were obtained in the same manner as the amplified assay with 40ms exposure time. Due to direct labeling, the fluorescent signal is generated from the biotin-streptavidin-Alexa Fluor® 568 complex, largely localized in the PEG layer. For imaging analysis, ROIs are defined as both the PEG layer and the internal droplet, the averaged signal from the ROI is used to represent the signal from that particle.

**Figure S1.**
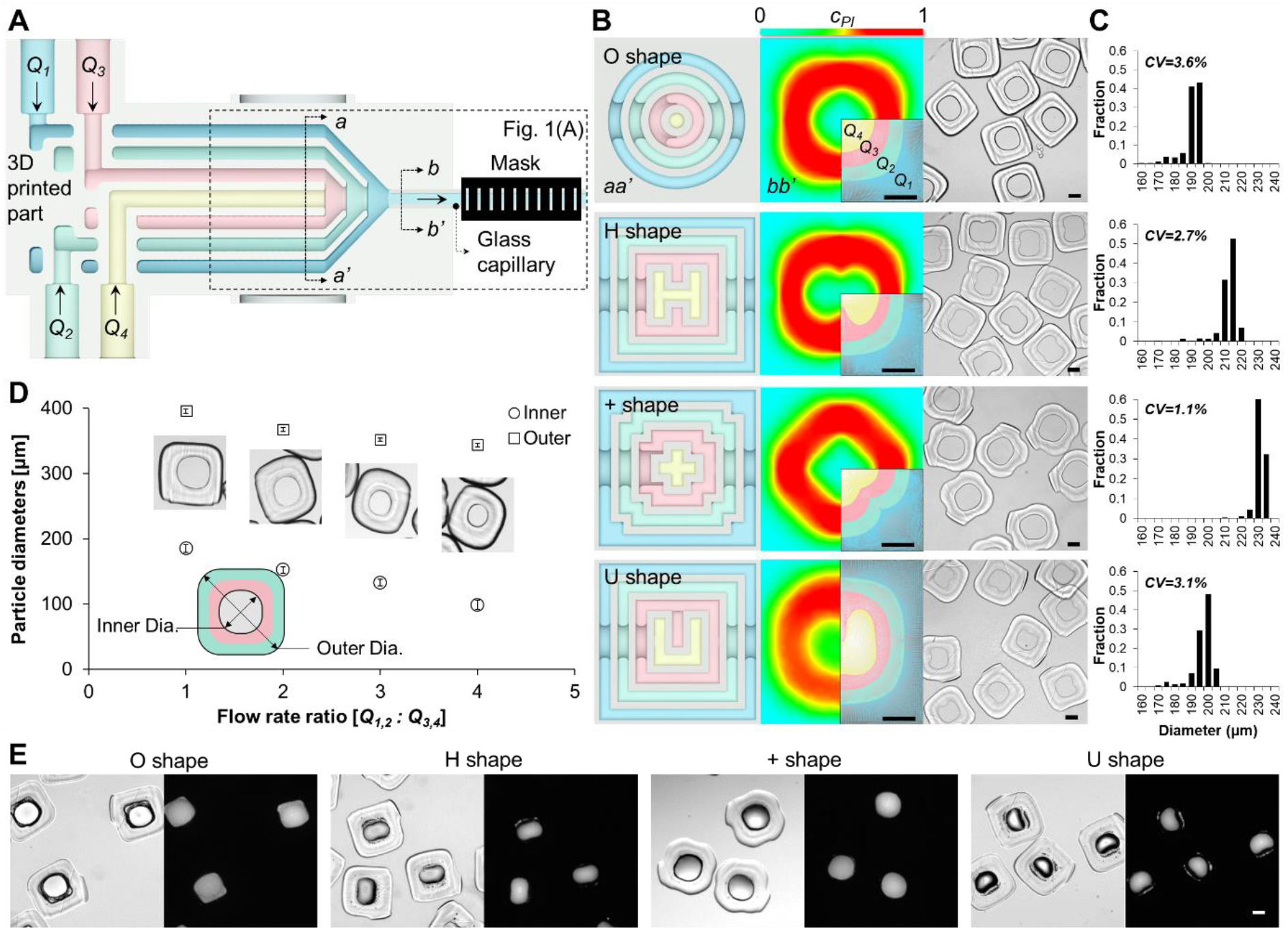
Shape-coded particles. **(A)** A cross-section of the 3D printed part joined with a square glass capillary is shown. The inner surfaces of the channels are colored blue, green, red and yellow to demonstrate they carry the four matching fluid streams (*Q*_*1-4*_) (see Figure 1A). **(B)** (left) The cross-section a-a’ of the device in (A) is shown for four different designs (O, H, Plus, and U shapes). (center) For the same designs, simulations of the concentration of PI (*c*_*PI*_) across the channel width at section b-b’ are shown for a laminar flow. The heat-maps show the normalized *c*_*PI*_ across the channel width when solving a convection diffusion equation. The insets show the location of the streamlines associated with each inlet flow *Q*_*1-4*_ within the square capillary cross-section (i.e. no diffusive blurring). (right) Manufactured particles corresponding to O, H, Plus and U-shape channel designs are shown. Scale bars are 100μm. **(C)** For each channel design, there is a narrow distribution in inner diameter of the cavities for manufactured particles. **(D)** Inner and outer diameters of O-shaped particles as depicted in the inset schematic are plotted against the flow rate ratios (*Q*_*1,2*_ : *Q*_*3,4*_). **(E)** Bright field and fluorescence images show FITC-containing aqueous droplets trapped within the cavities of the four different particle shapes (O, H, Plus, and U). Scale bar is 100μm.

**Figure S2.**
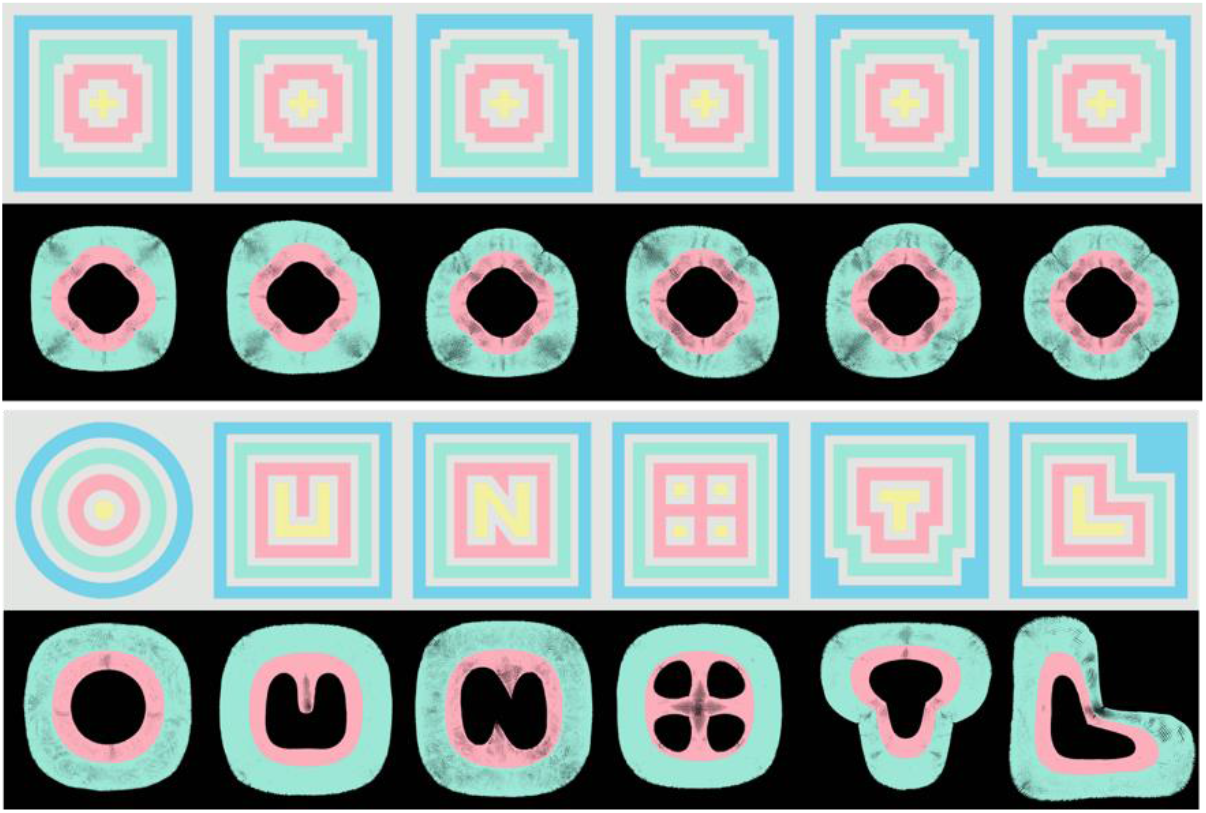
Shape coded 3D printed devices and simulated streamlines. Device cross-sections at a-a’ as indicated in Figure S1A and simulations of cross-sections of fluid streamlines corresponding to particles manufactured in Figure 2 are shown. The first and third row show the 3D printed device cross-sections and second and fourth row show the corresponding streamlines for flows through the device immediately above the simulation. The flow rate ratios for the simulations are *Q*_*1,2*_ : *Q*_*3,4*_ = 2:1. The cyan and magenta colored streamlines indicate the location of curable PPGDA and PEGDA streamlines respectively.

**Figure S3.**
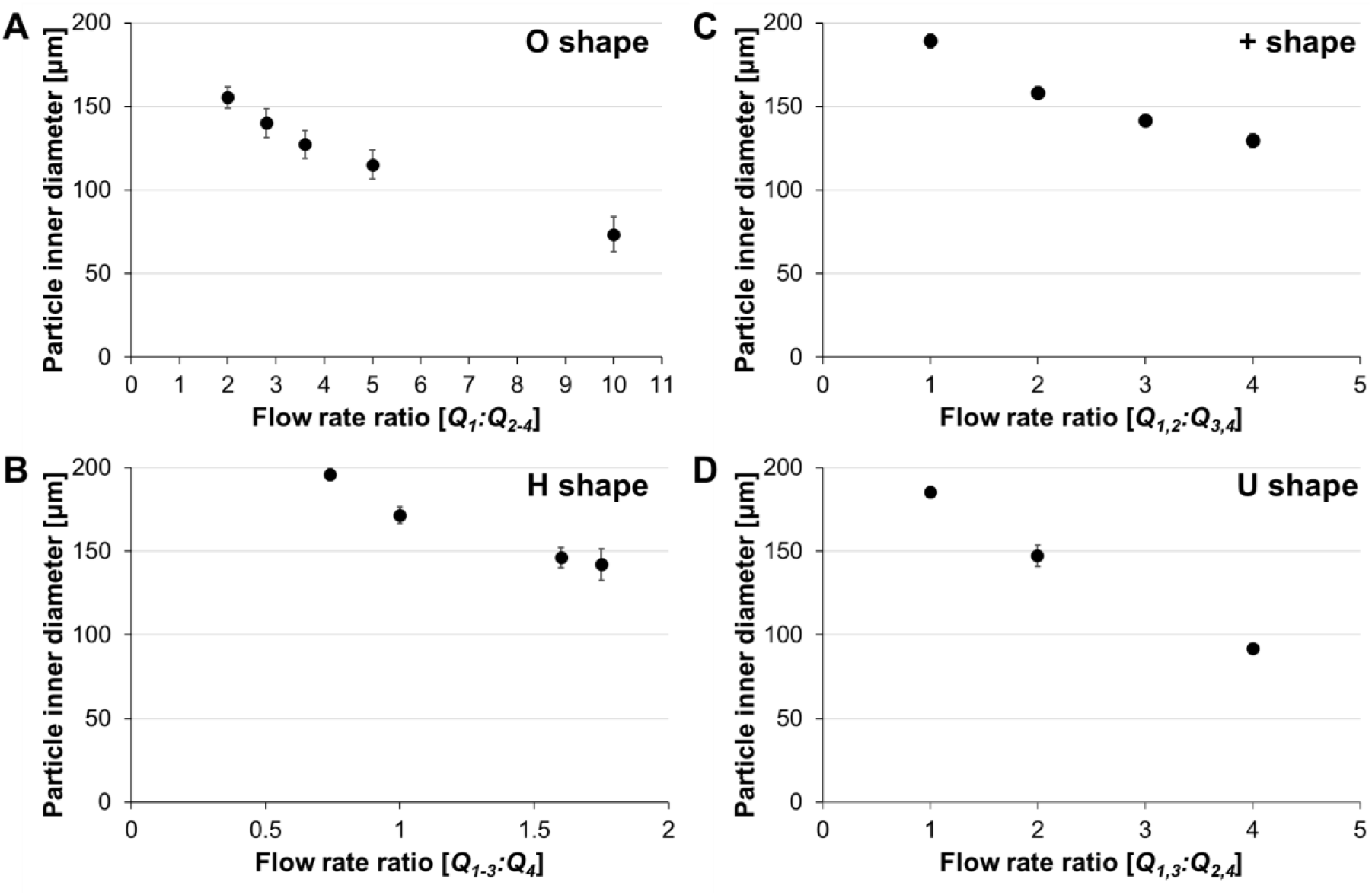
Modulation of particle inner diameter as a function of flow rate ratios for four different particle shapes. **(A)** The size of the O-shaped particle cavity decreased gradually as the flow rate ratio (*Q*_*1*_ : *Q*_*2-4*_) increased from 2 to 10. The experimental conditions are as follows: PI concentration of 5%, *τ*_*exp*_ = 0.3 s, *τ*_*d*_ = 1 s, *τ*_*s*_ = 5 s, and *Q*_*t*_ = 1.25, 1.45, 1.65, 1.6, 1.3 ml/min corresponding to each data point in the plot. **(B)** The size of the H-shaped particle cavity decreased gradually as the flow rate ratio (*Q*_*1-3*_ : *Q*_*4*_) increased from 0.74 to 1.75. The experimental conditions are as follows: PI concentration of 5%, *τ*_*exp*_ = 0.5 s, *τ*_*d*_ = 2.25 s, *τ*_*s*_ = 4 s, and *Q*_*t*_ = 1 ml/min for all the measurements. **(C)** The size of the plus-shaped particle cavity decreased gradually as the flow rate ratio (*Q*_*1,2*_ : *Q*_*3,4*_) increased from 1 to 4. The experimental conditions are as follows: PI concentration of 5%, *τ*_*exp*_ = 0.3 s, *τ*_*d*_ = 1, 1,25, 1.5, 1.75 s, *τ*_*s*_ = 4 s, and *Q*_*t*_ = 0.6, 0.9, 1.2, 1.5 ml/min, respectively, for the corresponding measurements. **(D)** The size of the U-shaped particle cavity decreased gradually as the flow rate ratio (*Q*_*1,3*_ : *Q*_*2,4*_) increased from 1 to 4. The experimental conditions are as follows: PI concentration of 2 and 4% in PPGDA and PEGDA, respectively, *τ*_*exp*_ = 0.5 s, *τ*_*d*_ = 1 s, *τ*_*s*_ = 5 s, and *Q*_*t*_ 1.2, 1.4, 1.6 ml/min, for the corresponding measurements.

**Figure S4.**
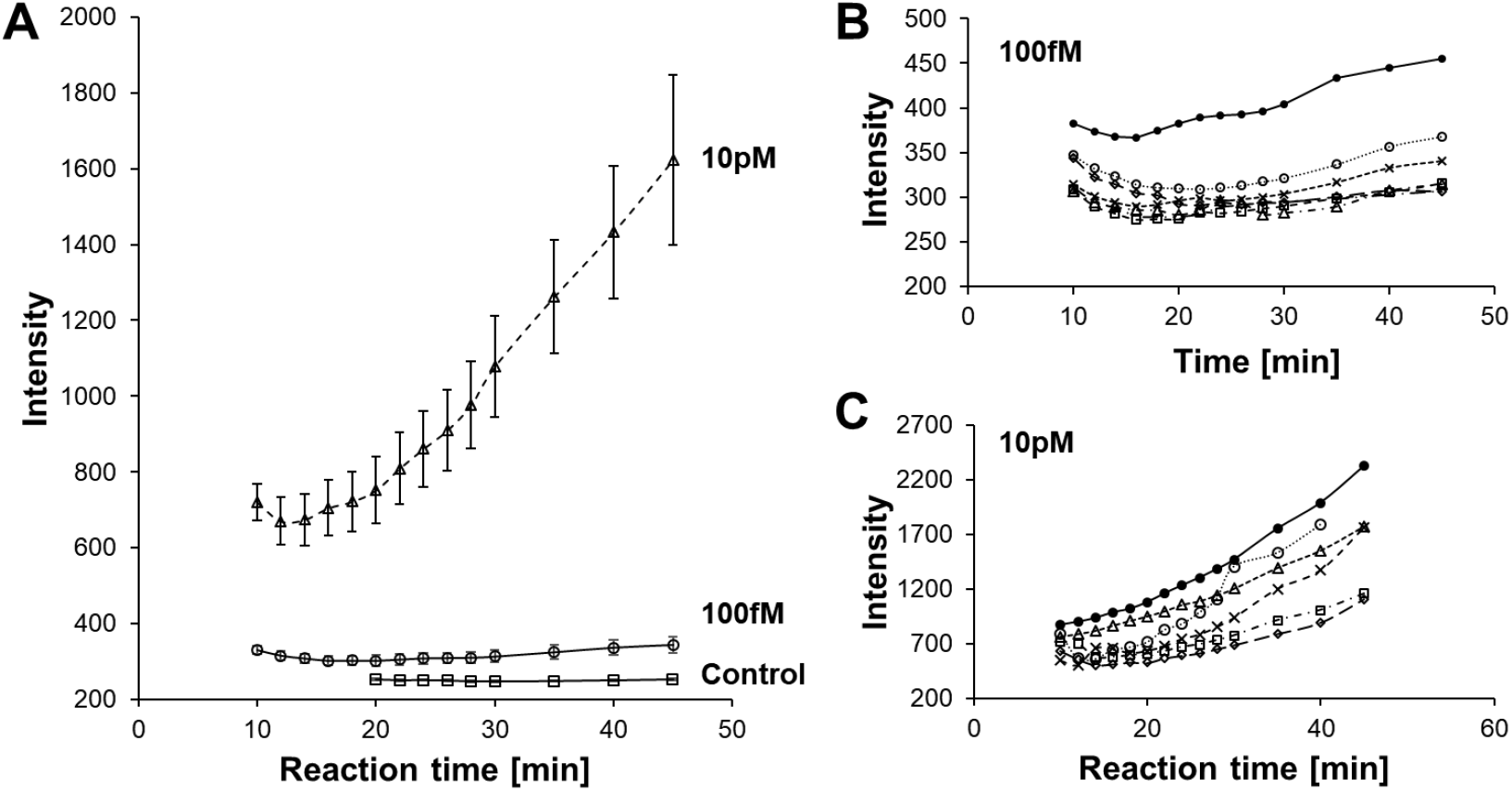
Time dependence of intensity change in dropicles using a QuantaRed assay. **(A)** Fluorescence signal increase over time from 10 to 45 min for binding of biotinylated particles with streptavidin-HRP at 10 pM and 100 fM concentrations. The control condition is the same particles and reagents without streptavidin-HRP. Data represents mean of intensity for dropicles with error bars representing standard error of the mean. Intensity drop at early time points is likely due to changes in shape/swelling upon introduction of the oil phase. (B-C) Fluorescence intensity from individual particles are shown over time for (B) 100 fM and (C) 10 pM concentrations of streptavidin-HRP respectively.

**Figure S5.**
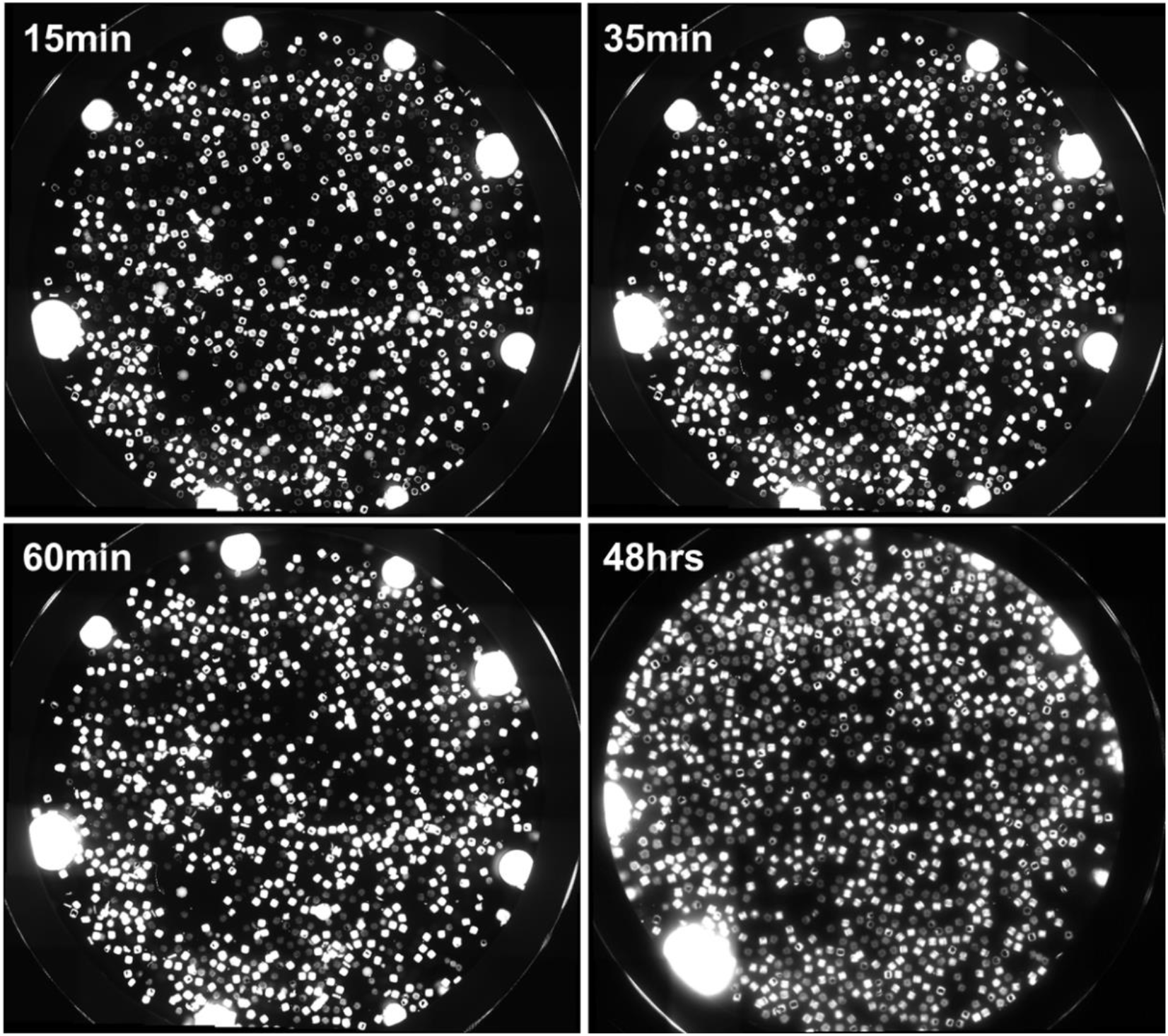
Minimal cross talk is observed between reactions in separate dropicles over time. Microscopic images showing the same well at 15 min, 35 min, 60 min, and 48 hrs after initiating the QuantaRed reaction. Two types of particles (with and without biotin) are introduced and incubated with 0.1 nM of streptavidin-HRP.

**Figure S6.**
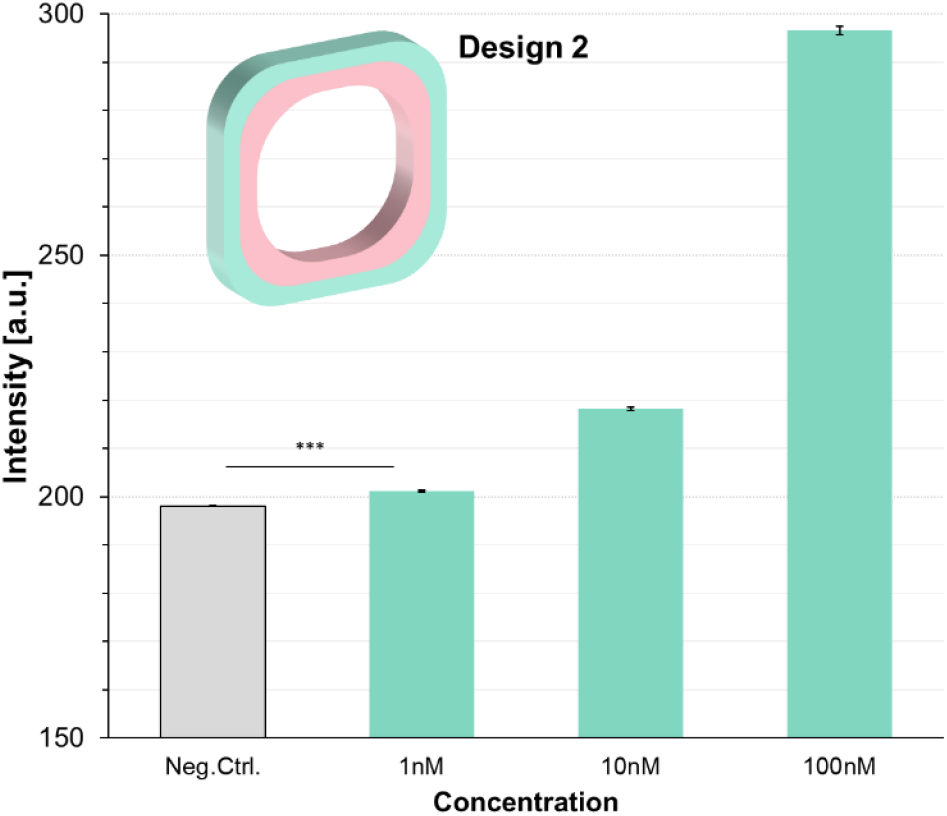
Direct binding assay using streptavidin. Intensity as a function of concentration for direct binding of fluorescent streptavidin without amplification using enzymes. Particles with design 2 are used. Error bars represent standard error. *** represents *p*<*0.001*.

**Figure S7.**
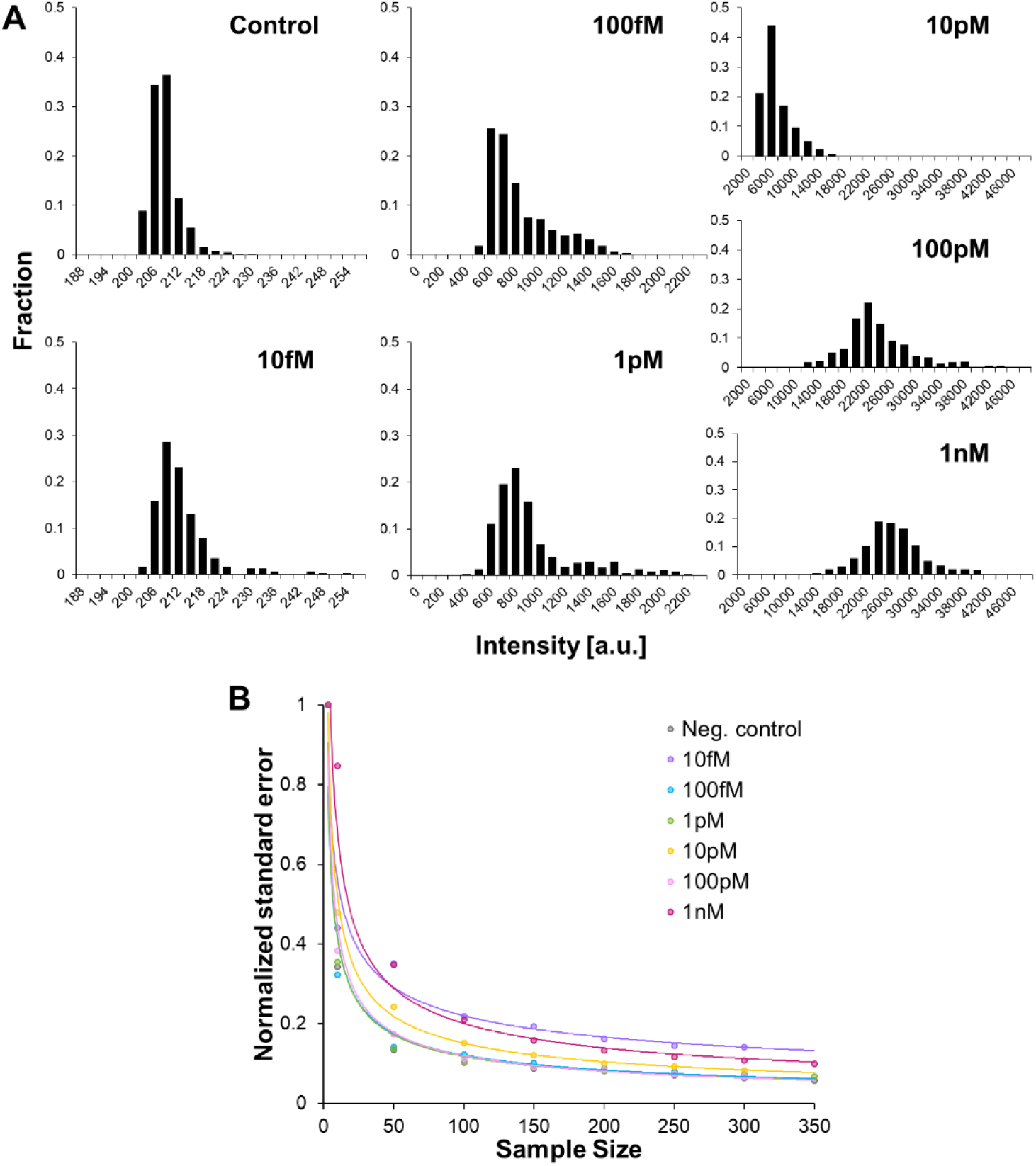
Swarm sensing with dropicles. (A) Histograms of fluorescence intensities for populations of dropicles formed from O-shaped particle design 2 after the QuantaRed assay readout at 45 min. Concentration of streptavidin-HRP ranges from 0 (control) to 1 nM. Results correlate with the mean intensity results reported in Figure 6C. (B) Normalized standard error of the mean vs number of dropicles in the sample for the amplified assay showing that a larger number of dropicles leads to a more accurate representation of the concentration of analyte.

**Table S1.** Comparison of multi-material particle fabrication techniques.

|  | Techniques<sup>(a)</sup> | CFL | CFL | CFL | SFL | HFL | HFL | SFL | MLL | MLL | IFS | HFT | 3D Co-flow |
| --- | --- | --- | --- | --- | --- | --- | --- | --- | --- | --- | --- | --- | --- |
|  | Year | 2006 | 2007 | 2007 | 2007 | 2010 | 2012 | 2014 | 2009, 2015 | 2012 | 2015-18 | 2018 | 2019 |
|  | Reference(s) | [20] | [22] | [21] | [25] | [29] | [32] | [7] | [33,34] | [40] | [36,37,39] | [35] | Current |
| Fabrication mechanism | Channel material<sup>(b)</sup> | PDMS | PDMS | PDMS | PDMS | PDMS | PDMS | PDMS | PDMS | PDMS | PDMS | COC | WS |
|  | Coating material<sup>(c)</sup> | × | × | × | × | × | NOA81 | PFPE | × | × | × | × | × |
|  | O<sub>2</sub> inhibition layer | ○ | ○ | ○ | ○ | ○ | × | ○ | ○ | ○ | ○ | N/A<sup>(d)</sup> | × |
|  | Inert flows | × | × | × | × | × | ○ | × | × | × | × | × | ○ |
|  | Hydrodynamic focusing | × | × | × | × | ○ | ○ | × | ○ | × | ○ | × | ○ |
|  | Organic solvents | × | × | × | × | × | ○ | ○ | × | × | × | × | ○<sup>(e)</sup> |
| Particle characterization | Particle size (~μm) | 3 | 100 | 180-270 | 1-6 | 50 | 50-200 | 20-250 | 100 | 200 | 100-500 | 50 | 100-200 |
|  | 3D shaped particles | × | × | × | × | × | × | × | ○ | ○ | ○ | ○ | ○ |
|  | On-demand size control | × | × | × | × | × | × | × | × | × | ○ | × | ○ |
|  | Amphiphilic particles | × | ○ | × | × | × | × | × | × | ○ | × | × | ○ |
|  | Uniform Droplet formation | × | ×<sup>(f)</sup> | × | × | × | × | × | × | ○<sup>(g)</sup> | × | × | ○ |

<sup>(a)</sup> CFL: Continuous Flow Lithography, SFL: Stop Flow Lithography, HFL: Hydrodynamic Flow Lithography, MLL: Maskless Lithography, IFS: Inertial Flow Sculpting, HFT: Hollow Fiber Template.
<sup>(b)</sup> PDMS: Polydimethylsiloxane, WS: WaterShed XC 11122, COC: Cyclic olefin copolymer.
<sup>(c)</sup> NOA81: Norland Optical Adhesive 81, PFPE: Perfluoropolyether.
<sup>(d)</sup> O<sub>2</sub> inhibition mechanism unknown.
<sup>(e)</sup> Data not shown.
<sup>(f)</sup> Droplet formed using multiple particles but lacked uniformity and controllability.
<sup>(g)</sup> Uniform volumes formed using multi-material particles, but throughput was limited by the multi-step fabrication process.

